# Curvature-guided chiral collective organization of myoblast tissues

**DOI:** 10.64898/2026.08.25.747076

**Authors:** Yuan Shen, Rushikesh Shinde, Wang Xi, Sushil Dubey, Yoann Le Toquin, Joanna Da Costa Oterelo Martins, Lucas Anger, Andreas Schoenit, Gianluca Grenci, Christophe Marcelle, René-Marc Mège, Raphael Voituriez, Andrew Callan-Jones, Benoit Ladoux

## Abstract

Surface curvature is a fundamental geometric cue in tissue morphogenesis, yet its role in guiding collective cell organization has remained elusive. Here, we show that curvature acts as a geometric control parameter that shapes supracellular alignment and chirality while modulating myogenic differentiation in myoblast tissues. Cells cultured on curved substrates self-organize into robust helical assemblies whose handedness is set, and can be reversed, by the sign of curvature: convex fibers produce right-handed helices, whereas concave channels invert the chirality. We identify a previously hidden clockwise bias in single-cell motion associated with the helical actin cytoskeleton. A minimal continuum theory coupling an effective chiral drive to curvature quantitatively captures the emergence and reversal of tissue-scale chiral alignment. On substrates with spatially varying curvature, local curvature gradients organize patterned multicellular architectures while preserving a global handedness. Curvature is also associated with myogenic state, with higher curvature linked to reduced or delayed differentiation. Together, these findings reveal how complex geometries shape the alignment, symmetry, and cellular state of living tissues.

**Significance statement:** Tissues develop on curved surfaces, yet it remains unclear how curvature shapes collective cell organization and function. We show that curvature can reliably control multicellular chirality, reversing the handedness of helical cell assemblies when surfaces switch from convex to concave. This geometric control extends to more complex curvature landscapes, which pattern distinct multicellular architectures. Curvature-driven order arises from coupling between local geometry and an intrinsic, actin-dependent clockwise bias in single-cell motion, revealing a route for chirality transfer across scales. Beyond organizing tissues, higher curvature is associated with reduced or delayed myogenic differentiation. Together, these results identify surface geometry as a design parameter for guiding tissue organization and influencing cellular state, with implications for morphogenesis and biomaterial design.

## Introduction

Tissues and organs in animal bodies adopt diverse three-dimensional shapes with complex surface curvatures that are intimately linked to their biological functions. The curved geometry of the cornea focuses light onto the retina, while the spiral cochlea sorts sound frequencies along its length; perturbing such geometries often leads to profound functional defects (*1, 2*). Surface curvature is therefore not merely a passive outcome of morphogenesis but an active physical constraint that shapes how tissues form and operate (*3*), underscoring the need to understand how geometry regulates collective cell organization and function (*4*). Yet systematically probing cell behavior on well-defined curved substrates has remained challenging, and direct observations of how cells sense and respond to curvature in three dimensions are still scarce (*5*).

Chirality, the property of a structure that cannot be superimposed on its mirror image, has been observed across biological scales (*6*). Intrinsic handedness can bias polarity in individual cells (*7*) and, in patterned cultures, can be amplified through cytoskeletal mechanics into coherent multicellular patterns (*8–11*). At the molecular and cellular scales, actin can self-organize into chiral spiral architectures (*12, 13*), whereas actomyosin-generated torques and formin-dependent cortical dynamics can drive chiral cortical flows (*14, 15*). The unconventional myosin 1D (Myo1D) provides a direct connection across biological scales, controlling organ laterality and transmitting handedness from actin-filament motion to cells, organs, and whole organisms (*16–18*). However, how these intrinsic chiral mechanisms interact with three-dimensional surface geometry remains poorly understood. A recent study showed that endothelial cells spontaneously form chiral helical arrangements in cylindrical channels (*19*). However, the study examined only cylinders with spatially uniform, same-sign curvature and therefore did not determine how changes in curvature sign or spatial variation influence cellular chirality. It remains unclear how local curvature couples to cell-intrinsic chirality and whether it can direct or even invert collective chiral order at the tissue scale. Addressing this gap is particularly important because developing organs, including the heart, gut, and liver, acquire left-right-asymmetric morphologies while adopting complex three-dimensional geometries (*20, 21*).

Here, we demonstrate that surface curvature governs not only the alignment of active tissues but also their handedness, serving as a geometric control parameter that organizes supracellular chirality and reverses its handedness between convex and concave surfaces. Myoblasts confined to cylindrical substrates self-organize into chiral supracellular structures whose handedness is set by curvature sign: convex fibers produce right-handed helices, while concave channels invert the chirality. Through live-cell imaging and quantitative analysis across multiple temporal resolutions, we reveal a subtle but robust clockwise bias in single-cell motion, which is associated with the helical actin cytoskeleton. A continuum theory coupling active chiral torque to local curvature quantitatively captures both the emergence and the reversal of tissue-scale chiral alignment. To probe how curvature gradients influence collective organization, we culture myoblasts on substrates with spatially varying curvature and find that they form patterned multicellular architectures, including coexisting nematically ordered and disordered domains, zig-zag nematic textures and spiral topological defects, that maintain robust right-handed order. Finally, we find that higher curvature is associated with reduced or delayed myogenic differentiation, linking geometric fields to collective organization, multiscale chirality transfer and cellular state.

## Results

### Helical superstructures on fibers and channels

To understand how surface curvature regulates collective dynamics, morphology and chirality of tissues, we cultured myoblasts on substrates of defined geometry. Myoblasts, spindle-shaped precursors of skeletal muscle fibers, naturally form nematic patterns on flat surfaces (*22, 23*). Here, we grew mouse C2C12 and immortalized human myoblasts (IHM) (*24*) on cylindrical PDMS fibers of varying diameters suspended in culture dishes (Fig. S1*A*). After 2-3 days, myoblasts self-organized into helical superstructures wherein cells were elongated and globally aligned with nematic order (Fig. 1*A*). The long axes of cells tilt relative to the fiber axis (*x*-axis) by an angle *θ* (negative for clockwise rotation about the normal vector ***N*** oriented from the substrate through the monolayer; see Fig. 1*I*), quantified by unwrapping the cylindrical surface to obtain the nematic director field ***n*** (Fig. 1*B*–*C*). We find that the magnitude of |*θ*| increases with fiber diameter *D* (Fig. 1*A*,*D*,*G*,*F*). The deviation of *θ* is large at *D* ∼ 800 μm due to the poor alignment and defect formation on large-diameter fibers (Fig. S2).

**Fig. 1.**
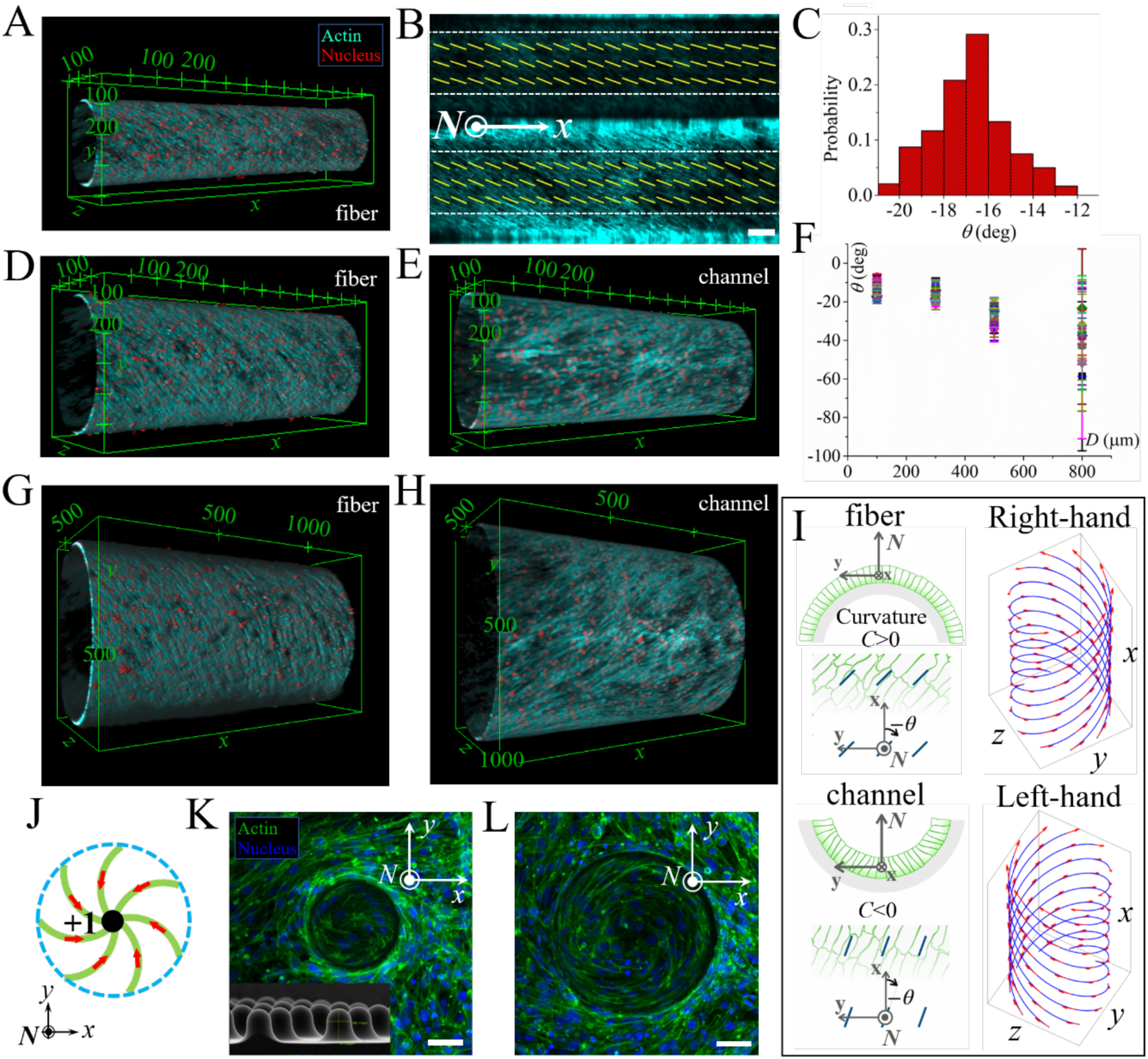
Helical superstructures of C2C12 myoblasts. (*A*) 3D visualization of a helical myoblast superstructure on a PDMS fiber with a diameter of 300 μm. (*B*) Myoblast layer obtained by unwinding the curved fiber (shown in *A*) and projected onto a flat surface (viewed from the outside of the fiber). The yellow short lines represent the director field, ***n***. Scale bar 100 μm. (*C*) Probability distribution of *θ* corresponding to the director field in (*B*). 3D visualization of myoblast helical superstructure on a fiber (*D*) and in a channel (*E*) both with diameters of 500 μm. (*F*) The tilt angle *θ* as a function of fiber diameter *D*. Each data point represents a single fiber. The error bars represent the standard deviation of the tilt angle of the cells on the fiber. 3D visualization of myoblast helical superstructure on a fiber (*G*) and in a channel (*H*) whose diameters are 800 μm and 1000 μm, respectively. (*I*) The schematic image showing the alignment of cells on fibers (top row) and channels (bottom row). The local normal ***N*** points from the substrate toward the apical side of the monolayer, and the tilt angle *θ* is measured relative to the cylinder axis. On both fibers and channels, the measured tilt has the same sign in this local convention, although its magnitude can differ. On a fiber, ***N*** points away from the cylinder center, whereas in a channel it points toward the center. Consequently, the same sign of local tilt produces opposite circumferential winding on the outside and inside surfaces, resulting in a right-handed helix on the fiber and a left-handed helix in the channel. (*J*) Schematic image showing a +1 topological defect with right-handed chirality. The *z*-stack projection of fluorescent microscopic image of C2C12 on a Gaussian-like dome (K) and a hemispherical dome (L). The inset shows the corresponding scanning electron microscopy (SEM) image of the dome. Scale bars 50 μm.

To exclude boundary and density effects, we varied fiber length *L* and culture time *t*. Neither parameter affects *θ* (Fig. S3), indicating that the tilt is an intrinsic geometric response rather than a confinement artefact. Strikingly, the director (***n***) consistently tilted in the same direction relative to the *x*-axis across independent experiments (Fig. 1*A*,*D*,*F,G*), indicating an intrinsic rather than spontaneous mirror-symmetry breaking. The helical pitch defined by this tilt corresponds to a right-handed superstructure (Fig. 1*I*).

To test the effect of curvature sign, we fabricated PDMS channels of comparable diameters and grew cells on their inner (concave) surfaces (Fig. 1*I*, S1*B*). In small-diameter channels, cell adhesion was less stable, and cells often detached and formed aggregates (Fig. S4). Such detachment was reported previously and was attributed to strong cell contractility (*25*). In larger channels, cells form stable helical superstructures. To verify that cell-matrix adhesion was maintained in stable concave geometries, we stained for paxillin, F-actin, and the fibronectin substrate coating. Paxillin-positive adhesions were present at the substrate-facing side in both channels and fibers, with similar organization (Fig. S5*A*-*E* and Fig. S6*A*-*E*). Cross-sectional confocal imaging confirmed that cells remained attached to the curved substrates and did not bridge the channel lumen (Fig. S5*F*-*I* and Fig. S6*F*-*I*). Thus, the fiber-to-channel transition preserves cell-substrate adhesion and basal-apical organization but changes the surface from convex to concave, thereby reversing the sign of curvature relative to the local basal-to-apical normal. Strikingly, reversing the local curvature experienced by cells from convex to concave reverses the chirality of the tissue superstructure: from right-handed on fibers to left-handed in channels (Fig. 1*E*,*H*,*I*, S7). Importantly, this reversal in global handedness does not require a reversal of the locally measured tilt angle. The tilt has the same sign on fibers and channels when defined relative to the local basal-to-apical normal, but it winds in opposite directions around the cylinder because the monolayer covers the outside of a fiber and the inside of a channel (Fig. 1I). Similar behaviors are also observed for IHM cells (Fig. S8).

To further confirm the chiral response of myoblasts, we grew cells on domes of different shapes (Fig. 1*J*-*L*; S9 for IHM). These domes introduce convex tips with non-zero Gaussian curvature, producing distinct supracellular organizations: cells form defects with a topological charge +1 at the tips. This behavior can be rationalized by our observation that myoblasts align along the circular interface between the domes and the flat substrate, thereby imposing an effective circular confinement with homogeneous alignment. Thus, the dome is topologically equivalent to a 2D disk with an Euler characteristic *χ* = 1, which implies a topological charge +1 (*26*). Importantly, all these +1 defects exhibit a spiral morphology with a robust right-handed chirality, echoing the helical patterns observed on convex fibers.

### Cell dynamics on fibers

To elucidate the dynamics underlying myoblast self-organization, we tracked single-cell motion of C2C12 cells (Fig. 2) and IHM cells (Fig. S10) on fibers. At early stages, actin filaments and cell trajectories appear disordered, but they progressively develop long-range alignment at later stages (Movie S1). To quantify this self-ordering process, we calculated the nematic order parameter, *S_n_*, of actin filaments and the velocity order parameter, *S_v_*, of cell motion (*Methods*). Here *S_v_* = 0 corresponds to random motion and *S_v_* = 1 to perfectly parallel or antiparallel velocities. We find both order parameters increase with time (Fig. 2*A* and S10*A*), confirming gradual self-ordering.

**Fig. 2.**
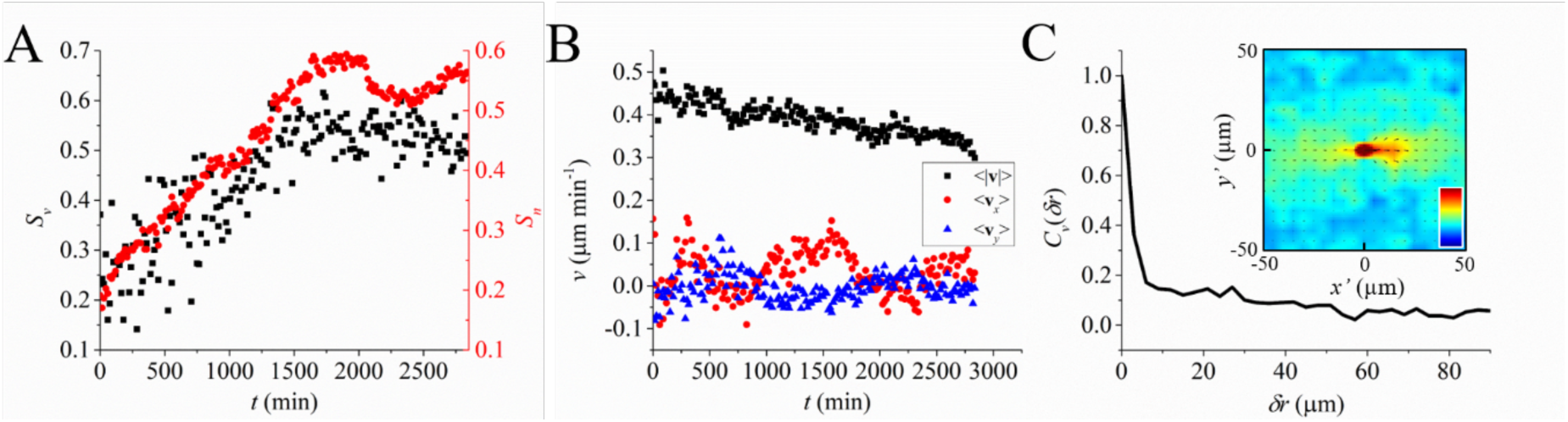
Dynamics of C2C12 on a fiber. (*A*) Temporal dependence of velocity order parameter (*S_v_*, black squares) and nematic order parameter (*S_n_*, red circles), respectively. (*B*) Temporal dependence of cell velocity. Black squares represent the mean value of cell speed. Red circles and blue triangles represent the mean values of the components of cell velocity along and perpendicular to the fiber axis, respectively. (*C*) 1D spatial velocity correlation function of C2C12 on the fiber. Inset: 2D spatial velocity correlation function of C2C12 on the fiber. The color bar scales linearly from -0.2 (dark blue) to 0.5 (dark red).

The mean cell speed (〈|***v***|〉) decreases over time as cell density increases (Fig. 2*B* and S10*B*). To assess whether cells collectively drift along a preferred direction, we examined the mean velocity components along the fiber’s longitudinal (*x*) and circumferential (*y*) axes. Both components fluctuate around 0 (Fig. 2*B* and S10*B*), indicating cells move bidirectionally and interact with each other through nematic (apolar) alignment.

We further calculated the equal time spatial velocity correlation function *C_v_*(*δr*) (*Methods*). Due to the absence of strong cell-cell adhesion, the correlations are short-ranged, decaying over about two cell lengths (Fig. 2*C* and S10*C*). When visualized in a local coordinate frame where each reference cell’s velocity defines the positive *_x_*′-axis, *C_v_* exhibits a distinct anisotropic pattern. This indicates cells tend to move in chains on fibers, following the ones ahead and leading those behind (insets in Fig. 2*C* and S10*C*).

### Chirality transfer from intracellular to multicellular scale

Knowing the dynamic self-organization process of helical superstructures on fibers, we next asked from where this supracellular chirality originates. Since individual myoblasts show no obvious morphological left-right asymmetry, we hypothesize that the handedness emerges from chiral cell dynamics.

We thus tracked the motion of isolated cells on flat glass substrates at low density over long durations (∼33 hours) (Fig. 3*A*). Cells exhibit short-range equal time spatial velocity correlations similar to those on fibers, with strong positive correlation along the direction of motion (Fig. 3*B*). The single cell (Lagrangian) temporal velocity correlation *C_v_*(*δt*) decays to 0 over a characteristic time of ∼200 min (Fig. 3*C*), interpreted as the cell persistence time. Although trajectories may seem qualitatively comparable to classical persistent random walks (Fig. 3*A*), comparing the unit tangent vector ***v***/*_ι_* to a given cell’s trajectory at different times might suggest chiral motion. Indeed, analysis of the velocity orientation cross product 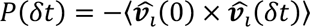, where the average is over initial times and cells, reveals a consistent positive bias for time lags *δt* shorter than the correlation time (Fig. 3*D*). This indicates a hidden but consistent clockwise rotational tendency of single-cell motion, consistent with chiral dynamics at the single cell scale. A recent study reported a counterclockwise rotation of nucleus within myoblast cells, but did not recognize this chiral single-cell motion probably due to the small time lag and the guiding effect of the grooved substrate they used (*27*).

**Fig. 3.**
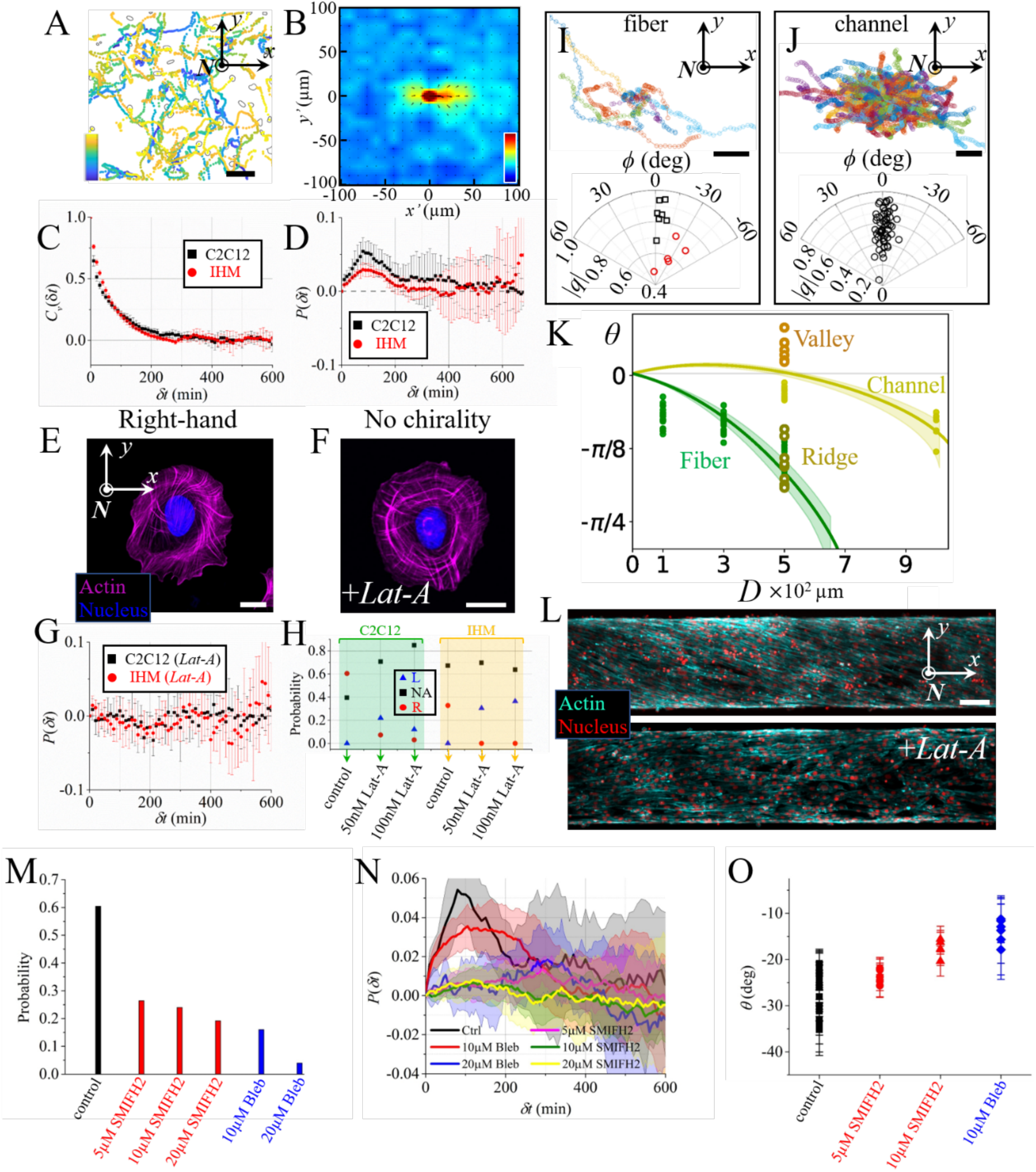
Chirality of myoblasts. (*A*) Trajectories of C2C12 on a flat glass substrate. The color bar represents the elapsed time which scales linearly from 0 (dark blue) to 2000 min (light yellow). Scale bar 100 μm. (*B*) 2D spatial velocity correlation function of C2C12 moving on flat glass substrates. The coordinate is rotated so that the velocity of the reference cells always points in the positive direction of the *x*’-axis. The color bar scales linearly from -0.2 (dark blue) to 0.5 (dark red). (*C*) Temporal velocity correlation functions of C2C12 and IHM on flat glass substrates. (*D*) The cross products of the unit velocity vectors of C2C12 and IHM as a function of time delay. Each data point in (*C*) and (*D*) is averaged over cells in six different fields of view (828 × 621 µm^2^). The error bars represent the standard deviation. Fluorescent microscopic images of a C2C12 cell (*E*) and a C2C12 cell treated with 50 nM *Lat-A* (*F*). Scale bars 20 μm. (*G*) The cross products of the unit velocity vectors of C2C12 and IHM as a function of time delay. Both C2C12 and IHM are processed with 50 nM *Lat-A*. Each data point is averaged over cells in five different fields of view (828 × 621 µm^2^). The error bars represent the standard deviation. (*H*) Statistics showing the probability of cells which form left (L, blue triangle) - or right (R, red circle) -handed spiral actin structure (NA, black squares, represents not applicable, i.e., actin cytoskeleton does not form clear spiral pattern as shown in (*F*)). Typical examples showing the trajectories of cells on a fiber (*I*, top) and in a channel (*J*, top). The starting points of the trajectories are shifted to the same point. Scale bars are 50 μm (*I*) and 100 μm (*J*), respectively. Distribution of the tilt angle, *ϕ*, of the trajectory structures of cells on fibers (I, bottom) and in channels (J, bottom) as a function of the anisotropy, |*q*|. The black squares and red circles in (*I*) represent cells on fibers of diameter 200 μm and 300 μm, respectively. The diameter of the channels in (*J*) is 500 μm. Each data point represents a single fiber or channel. (*K*) Data fit with the theoretical model. Scatter points are experimental data and the solid curves are model fits. (*L*) The *z*-stack projection of fluorescent microscopic images of C2C12 on fibers processed with (bottom) and without (top) *Lat-A*, respectively. scale bar 100 μm. (M) Statistics showing the probability of the C2C12 cells that form right-handed intracellular spiral actin organization at different conditions. About 50 random cells are measured in each condition. (N) The cross products of the unit velocity vectors of C2C12 cells at different conditions as a function of time delay. Each data point is averaged over cells in six different fields of view (828 × 621 µm^2^). The error bars represent the standard deviation. (O) The tilt angle *θ* of the nematic director field of helical supracellular structure on fibers at different conditions. Each data point represents a single fiber.

We then tracked cells on curved substrates at low seeding densities. When trajectories are shifted to a common origin, they form an anisotropic, tilted distribution (Fig. 3*I*–*J*). We quantify its anisotropy (|*q*|) and the tilt angle (*ϕ*) between its long axis and *x*-axis through the inertia tensor *M* (*Methods*) (*28, 29*). We note that |*q*| = 1 for a rod-like structure and |*q*| = 0 for a disk-like structure (*28*). On fibers, |*q*| is located between 0.4 and 1, and *ϕ* is always negative, consistent with clockwise cell tilt (Fig. 3*I*). In contrast, trajectories in channels are less anisotropic and exhibit only a weak negative bias (Fig. 3*J*), consistent with the small tilt angle *θ* of cell alignment in channels (Fig. S7). These results show that the chiral bias in single-cell motion emerges early and couples to the local substrate curvature to guide cell trajectories.

Based on these findings, we propose a theoretical model to disentangle the contributions of chirality and local substrate curvature in dense cell collectives and explain the formation of chiral supracellular structures on fibers and channels. We adopt a minimal description focusing on the dynamics of the director field ***n*** defined by the local cell orientation. This field is tangent to the substrate and can thus be parameterized by the mesoscopic alignment angle *θ* via 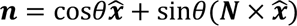. Here, we consider for simplicity an infinite cylindrical geometry mimicking fibers or channels (see Fig. 1*I*), where ***x***/ is the unit vector along the main axis; note that the model can be generalized to generic curved substrates. In the following we propose a systematic analysis of possible achiral- and chiral-nematic energetic couplings between *θ* and the substrate curvature to lowest orders in curvature. In the absence of flows, as is the case experimentally (Fig. 2*B*) and was also found with other spindle shaped cell types (*22*), our description is made minimally active by including the simplest active chiral term that reflects at the mesoscale the tendency of cells to rotate clockwise, as reported in Fig. 3*D*. The dynamics of *θ* at a point on the monolayer are thus described by

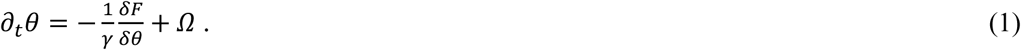

The first term on the right of this equation is the passive force conjugate to *θ*, where *F* is the Frank elastic energy which is specified below, and *γ* > 0 is the rotational viscosity (*26*). The active term in our model is the rotation rate *Ω*. A mathematically equivalent active chiral director-rotation term was previously introduced by Duclos et al. in a continuum model of confined cellular nematics (*22*). Such a term is allowed (*30*) when mirror symmetry is broken, detailed balance does not hold, and angular momentum is not conserved (because of the substrate). It represents the coarse-grained influence of microscopic chiral and active processes on the monolayer director. Its sign is motivated by the clockwise bias observed in single-cell motion (Fig. 3*D*), but its magnitude is not quantitatively derived from the single-cell measurements. Accordingly, our model treats *Ω* phenomenologically rather than deriving it from explicit cell-substrate and cell-cell interactions. For simplicity we assume that *Ω* is a constant material/substrate parameter, but in principle could depend on other quantities including the substrate curvature. The curvature dependence in our model is thus contained only in the passive, energetic part. In our minimal description the free energy density contains three terms that depend on the (mean) curvature, *C*, of the substrate-bound monolayer. The curvature is positive (negative) if the substrate bends away from (towards) ***N*** (Fig. 1*I*). For now we do not consider a gradient penalty in orientation since experimentally the tilt appears to be uniform on uniformly curved surfaces like channels and fibers, but we do so later (see “Order-disorder on conical substrates” below).

In the infinite cylindrical geometry relevant to our experiments with either fibers or channels, the energy density can be written:

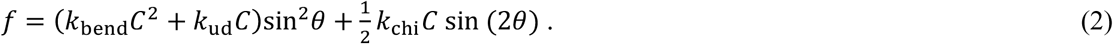

The first term on the right above is an anisotropic bending term, which is quadratic in curvature and has rigidity *k_bend_* > 0, favors alignment along /***x*** for large |*C*| (*31*). The second term is linear in curvature and achiral, and reflects a broken up-down symmetry in the adhered cells (*32*). Depending on the sign of *k_ud_C* this term favors alignment parallel or perpendicular to ***x***. The last term in Eq. (2) stems from chirality (*33*). Depending on the sign of *k_c_*_ℎ*i*_*C*, it favors alignment along the diagonals, *θ* = ±*π*/4. In a more mechanistic view, *k_ud_* could reflect, for example, an energy penalty of bending cells containing aligned stress fibers attached to the substrate, whereas the *k_c_*_ℎ*i*_ term could involve the chiral actin cytoskeleton, formin-dependent actin assembly, and myosin activity. However, our experiments do not identify a unique molecular origin for either coefficient. Moreover, an active curvature-dependent torque could produce the same steady-state angular dependence as the *k_c_*_ℎ*i*_ term in Eq. 3. (*34*).

Integrating the energy density over the surface, *F* = ∫ *fdS*, calculating its variational derivative with respect to *θ*, and restricting to steady-state conditions, we obtain from equations (1) and (2) the governing equation for the dependence of *θ* on curvature:

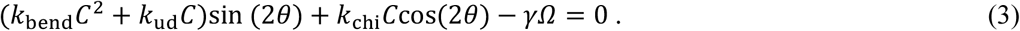

It predicts that myoblast alignment on substrates curved along one direction arises from the interplay between passive elasticity and the active rotation rate Ω. To test our model, we fitted the alignment data for C2C12 monolayers on fibers (*C* > 0) and channels (*C* < 0) of varying diameters, *D* = 2/|*C*|; see Fig. 1*F* and S7*A*. The fit, using the bootstrap method, is given in Fig. 3*K*. From the fit we estimated the non-dimensional active torque, *γΩ*/*k*_bend_ = (−6.46 ± 0.15) × 10^−5^*μ*m^−2^; and the ratio of energetic coefficients *k*_chi_/*k*_bend_ = (−1.68 ± 0.08) × 10^−3^*μ*m^−1^ and *k*_ud_/*k*_bend_ = (−1.21 ± 0.09) × 10^−3^*μ*m^−1^(see *SI* for details on the fit procedure). The negative value of the active rotation rate ***Ω*** is consistent with our measurements showing a net clockwise rotational bias of myoblasts (Fig. 3*D*). The negative value of *k*_ud_, combined with the negative ***Ω***, agrees with the observed stronger tilt on fibers than on channels (Figs. 1*F*, 3*I*-*J* and S7).

To investigate the intracellular origin of this chirality, we imaged the actin cytoskeleton of individual C2C12 cells at early time stages (∼10-20 min after seeding) at high resolution. Actin filaments frequently self-organized into spiral patterns that rotated around the substrate normal, ***N,*** while flowing inward from the cell periphery (Movie S2), reminiscent of actin dynamics previously reported in fibroblasts (*12, 13*). Among 48 randomly selected C2C12 cells, approximately 60% displayed these spiral actin patterns (Fig. 3*E*, *H*). We define this spiral organization of actin filaments as right-handed chirality: when the thumb points along the positive ***N*** direction, the curling of the other four fingers indicates the direction in which the actin filaments spiral inward. The remaining cells show no discernible spiral organization, and none displayed left-handed patterns (Fig. 3*H*, S11A).

We first tested whether these structures depended on actin polymerization. Treatment with Latrunculin-A (*Lat-A*) largely abolished the right-handed actin spirals (Fig. 3*F*). Among 41 treated cells, only ∼7% retain right-handed spirals, while ∼71% lose recognizable chirality and ∼22% develop left-handed spirals (Fig. 3*H*, S11B). Increasing *Lat-A* concentrations progressively reduces the occurrence of chiral spirals (Fig. 3*H*, S11C). Consistently, *Lat-A* treatment also eliminates the dynamic chirality of C2C12 cells: the velocity orientation cross product *P*(*δt*) fluctuates around zero across all timescales (Fig. 3*G*). At the multicellular level, *Lat-A* treatment suppresses the chirality of helical superstructures, resulting in a director field ***n*** aligned parallel to the *x*-axis (Fig. 3*L*). Interestingly, in IHM cells, the same treatment even reverses tissue handedness (Fig. S12). This inversion correlates with their intracellular actin organization: unlike C2C12, *Lat-A*-treated IHM cells display predominantly left-handed actin spirals (Fig. 3*H*, S13). Correspondingly, *P*(*δt*) for IHM cells shows a slight negative bias at short times (Fig. 3*G*).

Previous studies in *C. elegans* embryos showed that formin-dependent actomyosin dynamics can generate chiral cortical flows and torques (*14, 15*). Motivated by these findings and by the inward rotational actin flows observed in myoblasts (Movie S2), we tested whether chirality also depends on formin-sensitive actin assembly and myosin-II activity using SMIFH2 and blebbistatin, respectively. SMIFH2 reduced intracellular actin spirals and weakened the clockwise bias in single-cell motion in a concentration-dependent manner (Fig. 3*M*, *N*). At low concentrations, treated cells still formed helical structures on fibers, but the absolute tilt angle |*θ*| decreased with increasing concentration (Fig. 3*O* and S14). At 20 μM, cell death and incomplete monolayer formation prevented reliable tissue-scale analysis (Fig. S14). These observations suggest that formin-sensitive actin assembly contributes both to intracellular actin chirality and to its transmission from the intracellular and single-cell scales to supracellular organization.

Blebbistatin similarly reduced the fraction of cells displaying spiral actin organization (Fig. 3*M*). At low concentration (10 μM), cells retained a clockwise bias in single-cell motion (Fig. 3*N*) and still formed helical supracellular structures on fibers (Fig. S14), although with a smaller absolute tilt angle |*θ*| (Fig. 3*O*). By contrast, at higher concentration (20 μM), cells lost the detectable clockwise bias in single-cell motion (Fig. 3*N*) and failed to organize into helical supracellular structures (Fig. S14). These findings are consistent with a role for myosin II activity in maintaining chiral cell dynamics and in transmitting chirality from the intracellular and single-cell scales to the tissue scale. However, because strong myosin II inhibition can also affect cell contractility, motility, and monolayer organization more broadly, we interpret these results as supporting, rather than proving, a causal role for actomyosin activity in multiscale chirality transfer.

Together, these results link intracellular actin spirals and clockwise-biased cell motion to curvature-guided helical tissue organization. *Lat-A* indicates that actin polymerization is required, while SMIFH2 and blebbistatin implicate formin-sensitive actin assembly and myosin-II activity in transmitting chirality across scales.

Finally, we asked whether single-cell chirality could also be detected within dense monolayers. In dense monolayers, spiral actin patterns were transient and rapidly replaced by local nematic alignment (Movie S3). We therefore used centrosome positioning as an alternative readout of single-cell chirality, following the approach of Zhang et al. (*19*), and adapting it to the nematic symmetry of our system (*Methods*). The nucleus-centrosome axis showed a weak clockwise bias relative to the local nematic director (Fig. S15). Thus, although chirality is expressed differently in isolated cells and dense monolayers, likely because of cell-cell interactions and collective nematic alignment constraints, cells within the tissue retain a handed bias consistent with their chiral dynamics and tissue-scale organization.

### Order-to-disorder on conical substrates

We previously found that cells on cylindrical fibers develop a uniformly tilted nematic organization, with the tilt angle increasing with fiber diameter. To investigate how spatial variations in curvature affect collective alignment, we cultured C2C12 myoblasts on a truncated conical PDMS substrate, whose diameter increases smoothly along its symmetry axis. Near the tapered end, cells formed a uniformly tilted monolayer (Fig. 4*A*-*B*), resembling the organization observed on cylindrical fibers. With increasing local diameter, however, nematic order progressively weakened. This loss of order was accompanied by the emergence of +1/2 and -1/2 topological defects (Fig. 4*A*), regions where the nematic order parameter, *S_n_*, decreased to close to zero (Fig. 4*C*).

**Fig. 4.**
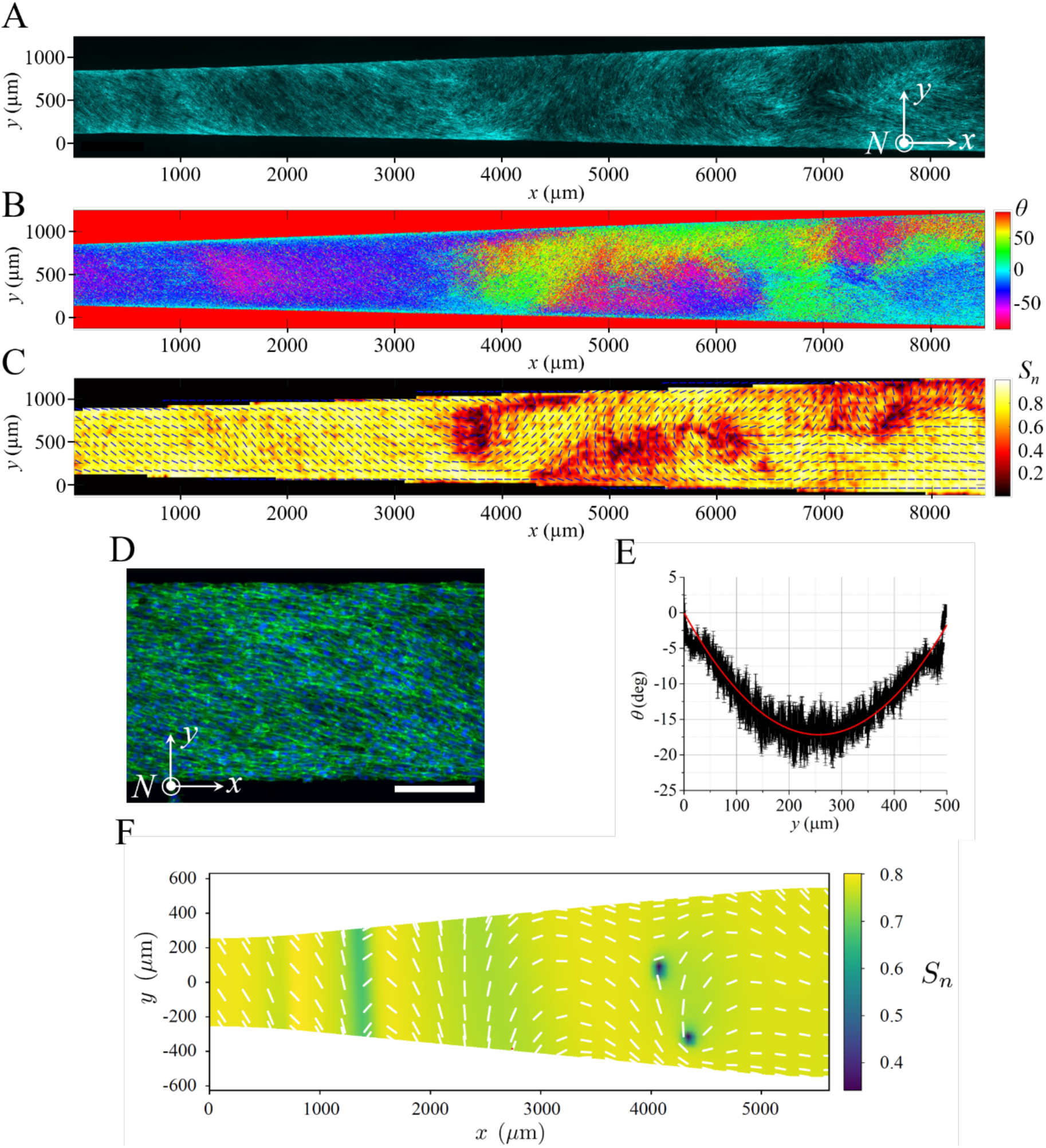
Transition from an ordered to disordered chiral nematic of C2C12 myoblasts on a conical surface. (A) The z-stack projection of fluorescent microscopic image of the actin structure of C2C12 on a truncated cone. (B) Color map showing the cell tilt angle, *θ*, in degrees. (C) Nematic order of actin structures revealed by order parameter *S_n_* and the corresponding director field (black bars). Topological defects coincide with dark spots (regions of small values of *S_n_*) in the heat map. (D) Cell monolayer confined to an adhesive strip on a flat substrate. Scale bar is 200 *μ*m. (E) Tilt profile in the *y*-direction for the monolayer in D. A fit to *θ*(*y*) using Eq. 3 (augmented by the gradient penalty *K∇*^2^*θ* and neglecting curvature terms) yields *γΩ*/*K* ≈ 2.6 × 10^−4^*μm*^−2^. (F) Simulation of the nematic tensor ***Q*** (see *SI* for parameter values).

To explain the coexistence of ordered and disordered regions, we extended our theoretical framework to allow both the tilt angle, *θ*, and the magnitude of nematic order, *S_n_*, to vary in space. We reformulated the model in terms of the nematic order tensor ***Q*** = *S_n_*(***nn*** − ***I***/2), with ***I*** the two-dimensional identity tensor, whose dynamics is ruled by energy minimization (gradient flow) and an active rotation ***Ω*** (see *SI* for details). The coupling between ***Q*** and surface curvature is analogous to the curvature coupling previously introduced for the director ***n***. The model also includes an energetic cost associated with spatial variations in nematic alignment. In the strongly ordered limit, this contribution reduces to the one-constant Frank elastic energy, *K*(*∇θ*)^2^/2, where the Frank constant *K>0*, whose effects are illustrated in a simple geometry (Fig. 4*D*-*E*). Using this extended model, we simulated the emergence of nematic order on a truncated cone from disordered initial conditions that mimic experimental seeding. Consistent with our experiments, the nematic field near the narrow, highly curved end rapidly reached a steady state characterized by a uniform tilt. Toward the wider end, where curvature is lower, alignment was weaker and evolved more slowly, resulting in persistent disordered structures, walls, and topological defects (Fig. 4*F*). Overall, we find that a tapered substrate, mimicking structures such as the tips of epithelial tubes (*35*), leads to spatially segregated tilted ordered and disordered multicellular domains, suggesting that curvature can act as a geometric field that tunes the degree and spatial variation of collective alignment in monolayers.

### Zig-zag nematic structures on wavy substrates

We have shown that cells form right- and left-handed helical superstructures on fibers and channels, geometries characterized by uniform curvatures of opposite sign. To determine whether such chiral organization persists on more complex geometries featuring curvature sign changes, we next examined a single substrate exhibiting spatially varying curvature with both concave and convex features.

C2C12 myoblasts cultured on PDMS substrates with a periodic wavy topography (Fig. S1*6A*) spontaneously organize into a zig-zag nematic pattern, forming robust right-handed superstructures irrespective of the local curvature sign (Fig. 5*A*,*B*; Fig. S1*6B* for IHM cells). On ridges, cells tilt clockwise relative to the zero-curvature (*x*) axis, consistent with the behavior on convex fibers, making a right-handed superstructure. Strikingly, in valleys whose local geometry mimics concave channels, cells tilt counterclockwise and thus produce an overall right-handed superstructure, opposite to the left-handed configuration observed in isolated channels.

**Fig. 5.**
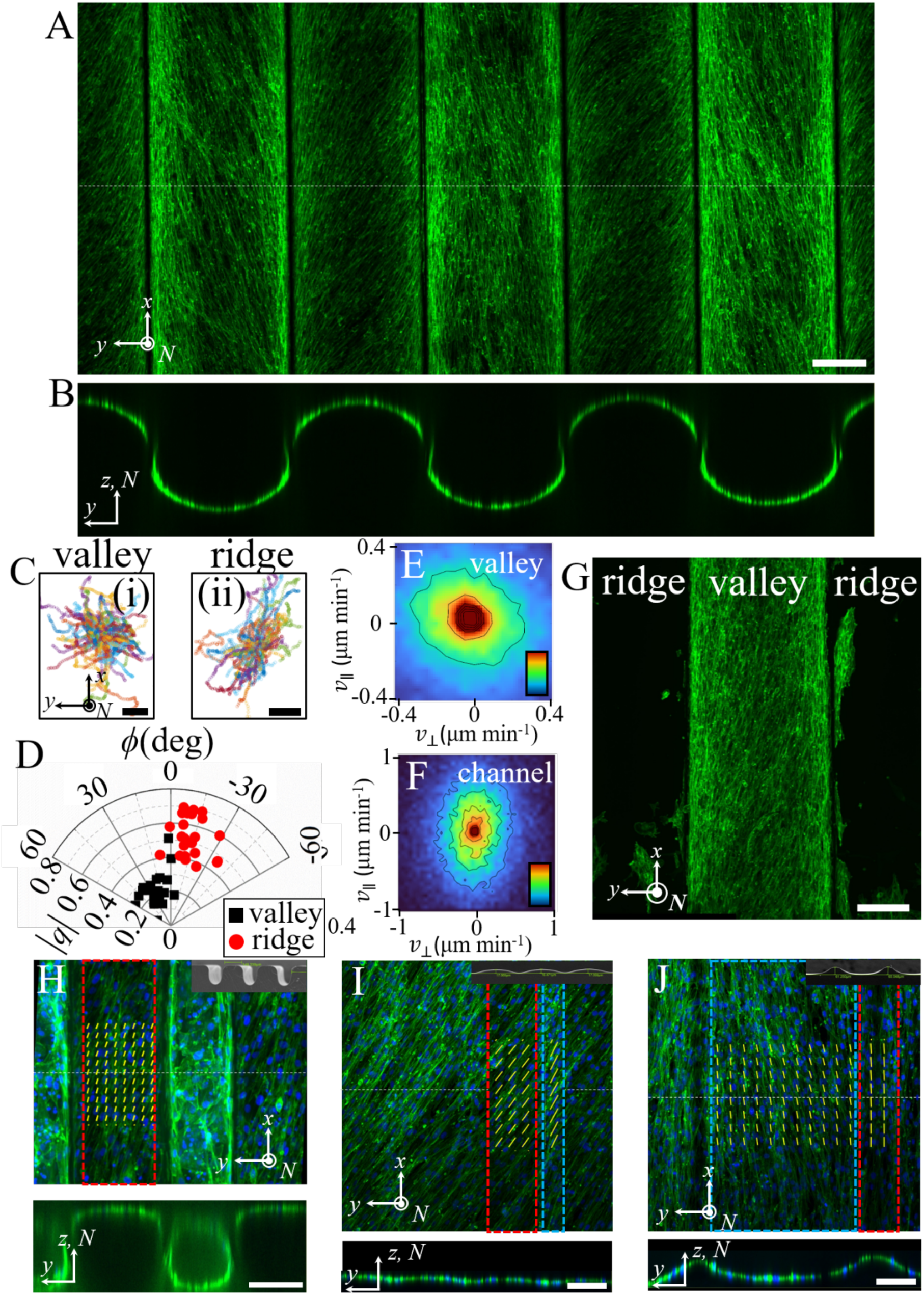
C2C12 on wavy substrates. (*A*) The *z*-stack projection of fluorescent microscopic image of the actin structure of C2C12 on a wavy substrate (apical view). Scale bar 200 μm. (*B*) The cross section of the sample in the *yz*-plane. (*C*) Cell trajectories which are shifted so that they start at the same position ((i): valley; (ii): ridge). Scale bars 100 μm. (*D*) Polar plot of the tilt angle, *ϕ* (with respect to the *x*-axis), of the trajectory structures of cells on ridges (red) and in valleys (black) as a function of the anisotropy, |*q*|. Each data point represents a single ridge or valley. (*E*-*F*) Histogram of cell velocities in valleys (*D* = 500 μm) (*E*) and in channels (*D* = 500 μm) (*F*. *v*_||_) and *v*_⟂_ represent the velocity component along and perpendicular to the principal direction of zero curvature. The histograms are computed over 5 different valleys and channels (through a tracking time period about 20 hours). The color bar scales linearly from 0 (dark blue) to 200 (dark red). (*G*) Fluorescent microscopy image of the actin structure of C2C12 on a wavy substrate in which the ridges are processed with pluronic to prevent cell attachment. Scale bar 200 μm. (*H*-*J*) The *z*-stack projections (top) and the cross sections (bottom) of the fluorescent microscopic images of C2C12 (green: actin; blue: nucleus) on different wavy substrates (apical view). The red and blue dashed rectangles represent regions of ridges and valleys, respectively. The insets show the corresponding SEM images of the PDMS substrates. The yellow short lines represent the director field. Scale bars 100 μm.

Theoretical fits reproduce the ridge alignment but fail to account for the positive tilt (*θ* > 0) observed in valleys (Fig. 3*K*). This behavior contrasts with that in channels, which also exhibit negative curvature but where cells tilt clockwise (Fig. 1*I*). Simultaneously, single-cell tracking shows that cell trajectories on ridges form anisotropic bundles with negative tilt angles *ϕ*, closely resembling those observed on fibers (Fig. 5*C*-*D*). However, trajectories in valleys differ markedly from those in channels: cells in valleys preferentially migrate perpendicular to the zero-curvature axis and exhibit positive tilt angles, whereas cells in channels migrate predominantly parallel to this axis and exhibit negative tilt angles (Fig. 5*C*-*F*; Movies S4-S5). Despite having similar curvature magnitude and sign, valleys and channels differ in two key aspects: (i) curvature in valleys is spatially nonuniform and vanishes at the boundaries adjoining ridges, imposing (ii) distinct boundary conditions. To isolate the role of (i) and (ii), we cultured cells on wavy substrates in which only the valley region was rendered adherent. Remarkably, the tilt then reversed sign, matching the alignment observed in channels (Fig. 5*G*). This result suggests that boundary conditions imposed by adjacent ridges and possibly local curvature gradients govern the alignment of cells within valleys.

We further tested curvature dependence using wavy substrates of more complex geometries (Fig. 5*H*– *J*; Fig. S1*6C* for IHM cells). On ridges with positive curvature, cells consistently form right-handed superstructures similar to those on fibers. In valleys, the behavior varied with curvature magnitude: strong concavity impairs cell spreading (Fig. 5*H*), weak concavity yields nearly uniform alignment (Fig. 5*I*), and intermediate concavity produces a small counterclockwise tilt *θ*, akin to that in Fig. 5*A* (Fig. 5*J*). These findings highlight that the coordinated interplay between local curvature gradients and boundary conditions actively directs collective cell organization, revealing curvature as a subtle cue for spatially programming cellular alignment.

### Curvature dependence of myogenic differentiation

To determine whether substrate curvature influences myogenic differentiation, we cultured C2C12 and IHM myoblasts on cylindrical fibers of different diameters. After the helical monolayers had formed, cells were transferred to differentiation medium, where they progressively fused into multinucleated myotubes (Fig. 6*A*). We first assessed nuclear p27, a cyclin-dependent kinase inhibitor whose expression increases during C2C12 differentiation and contributes to differentiation-associated cell-cycle withdrawal and myogenic progression (*36, 37*). Although p27 is not a muscle-specific differentiation marker, increased nuclear p27 provides a useful readout of differentiation-associated cell-cycle exit in this context. In both C2C12 and IHM cells, the fraction of p27-positive nuclei increased with fiber diameter (Fig. 6*B*–*C*; Figs. S17– S18). Thus, cells on wider, less strongly curved fibers showed greater differentiation-associated cell-cycle withdrawal.

**Fig. 6.**
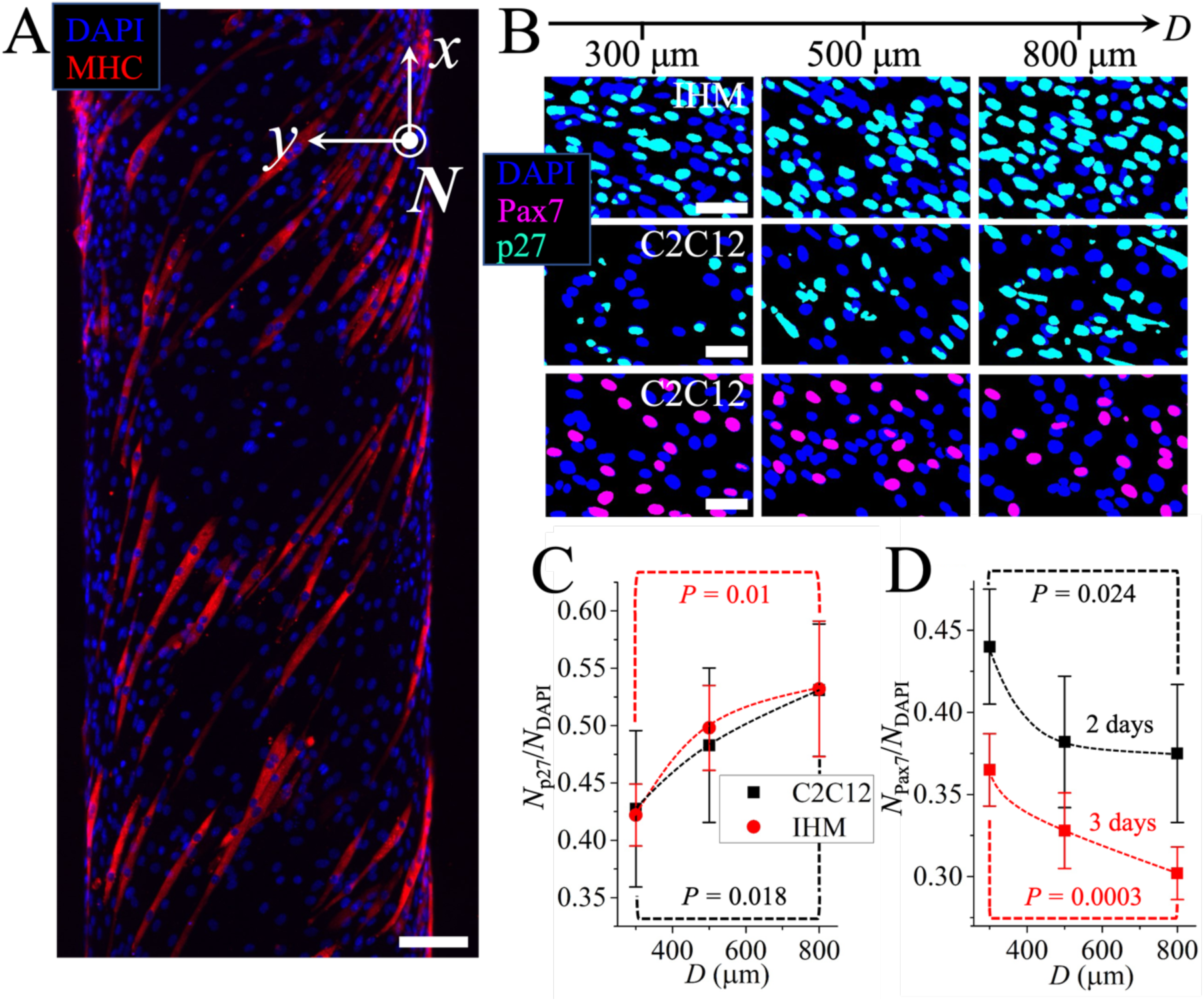
Cell differentiation on fibers. (*A*) The *z*-stack projection of fluorescent microscopic image of C2C12 cells on a fiber. Scale bar 100 μm. The blue and red signals represent the DAPI and myosin heavy chain (MHC). (*B*) Nuclei segmentation of the immunostaining images of IHM (top row) and C2C12 (the middle and the bottom rows) as a function of fiber diameter *D*. Blue: DAPI; magenta: Pax7; cyan: p27. Scale bars 50 μm. Full field-of-view images are provided in Fig. S17-S19. (*C*) Quantification of the ratio between the number of cells that express marker (p27-positive) and the total number of cells as a function of fiber diameter. (*D*) Quantification of the ratio between the number of C2C12 cells that express marker (Pax7-positive) and the total number of C2C12 cells as a function of fiber diameter. (black squares: 2 days in differentiation media; red circles: 3 days). The indicated *P*-values in (*C*) and (*D*) compare the 300- and 800- µm fiber groups using two-sided Welch’s t-tests. Each data point is averaged over 6 different fibers. The error bars represent the standard deviation. The dashed lines are guides to the eye.

We obtained complementary evidence in C2C12 cells by staining for Pax7, a marker of myogenic precursor states that is downregulated as myoblasts differentiate (*38, 39*). The fraction of Pax7-positive nuclei decreased over time and, at a given time point, decreased with increasing fiber diameter (Fig. 6*B*, *D*; Fig. S19). Pax7 staining was not sufficiently robust in IHM cells under our experimental conditions for quantitative analysis. Together, the increase in the p27-positive fraction and decrease in the Pax7-positive fraction are consistent with enhanced myogenic progression on wider fibers, supporting the conclusion that stronger curvature delays differentiation.

## Discussion and conclusion

Although cells have long been known to sense surface curvature (*40, 41*), its role in shaping tissues has remained underexplored until recent advances in microfabrication, imaging, and theoretical modeling (*3, 42*). Emerging work has shown that curvature can guide collective migration of epithelial (*43–47*) and human bone marrow stromal cells (*48*), modulate pre-osteoblast organization (*49*), affect cell focal adhesions, nuclear shape, and gene expression (*47*), regulate epithelial monolayer thickness (*50*) and promote nematic alignment of myoblasts (*51*), endothelial (*19, 52*) and smooth muscle cells (*53*). A theoretical “minimal cell” model that reproduced cell motion on different curved surfaces was also proposed recently (*54*). Yet, most studies have focused on fixed samples or a single geometry (*19, 43–45, 51, 52*), leaving open how local curvature and curvature gradients dynamically interact with intrinsic cellular properties to regulate tissue-scale organization. Meanwhile, intrinsic cellular chirality has been proposed as a potential origin of tissue-scale left-right asymmetry (*8, 10, 17, 19*). Yet how such single-cell chirality couples to geometric cues and scales up to robust, collective chiral order has remained unclear. In particular, whether curvature merely constrains emergent chiral patterns or actively selects and controls their handedness has not been established.

In this work, we systematically map how myoblast morphology and dynamics depend on curvature, ranging from convex fibers to concave channels, truncated cones, wavy substrates, and surfaces with non-zero Gaussian curvature. We uncover a previously unrecognized clockwise bias in single-cell motion and show, using a continuum model, that the interplay between curvature-dependent alignment and an effective active rotation rate produces macroscopic supracellular chiral order. Our work on emergent chirality stemming from cell-substrate interaction is distinct from earlier work where large scale chirality emerges from active torques within the material, independently of a substrate (*14*). This curvature-chirality coupling produces a hierarchy of chiral superstructures and persists even on substrates with spatially varying curvature, where curvature gradients guide the formation of coexisting nematically ordered and disordered domains, zig-zag nematic textures and spiral topological defects. Related curvature-coupled collective behaviors have also been reported in endothelial and epithelial tubes, although their dynamical manifestations and dependence on cell-cell adhesion differ. In endothelial microvessels, Zhang et al. observed robust right-handed helical cell alignment, linked tissue handedness to cellular chirality measured *in situ* from organelle positioning, and showed that junctional perturbations and tissue fluidity modulate the strength of the helical order (*19*). In MDCK cylindrical epithelia, Glentis et al. instead observed persistent circumferential collective rotation in either the clockwise or counterclockwise direction, rather than a fixed handed alignment (*43*); this coordinated rotation required stable cadherin-mediated cell-cell junctions. By contrast, myoblasts exhibit comparatively weak cell-cell adhesion and short-ranged velocity correlations, yet form a robust handed nematic tilt. Together, these observations suggest that strong cadherin-mediated cohesion may not be strictly required for curvature-guided handed alignment, but can regulate the spatial transmission, coherence, and dynamical expression of cellular chirality in different tissue types.

Beyond collective organization, our results suggest that surface curvature modulates myogenic differentiation, with higher curvature associated with reduced or delayed differentiation. Because thinner fibers support stronger nematic alignment while wider fibers provide weaker alignment, these findings suggest that curvature-guided tissue organization is coupled to cellular state: high curvature stabilizes aligned helical organization while delaying differentiation, whereas lower curvature is associated with weaker alignment and enhanced differentiation. We previously reported that topological defects trigger high compressive stress and enhance myoblast differentiation (*55*). In the present system, low-curvature fibers, corresponding to larger diameters, display poorer global alignment and more frequent defects than highly curved fibers (Fig. 1*F*; Fig. S2). These observations raise the possibility that curvature-dependent changes in nematic order and defect formation may influence the differentiation response. Since defects and orientational disorder can be associated with local mechanical and dynamical heterogeneity (*56*), we asked whether fibers of different diameters also exhibit different levels of cell-motion heterogeneity. To this end, we quantified spatiotemporal fluctuations in cell velocity on fibers of different diameters (Figs. S20-S21; *Methods*). Although speed fluctuations did not show a clear diameter-dependent trend, the transverse component of cell-velocity fluctuations was substantially larger on fibers of larger diameter (Fig. S20*I*; Fig. S21*E*). This result suggests that low-curvature, poorly aligned tissues exhibit enhanced transverse dynamical heterogeneity, which may be related to the increased differentiation observed on larger fibers. However, because velocity fluctuations do not directly measure mechanical stress, future measurements of traction forces and intercellular stresses on curved surfaces will be required to determine whether curvature-dependent mechanical stress fluctuations contribute to myogenic differentiation.

Together, our results identify curvature as a physical morphogenetic cue that shapes collective tissue organization and is associated with myogenic state. Controlling local curvature and its gradients may therefore offer a means to guide collective behavior and influence differentiation-associated cellular states in engineered tissues. More broadly, our work provides a physical framework linking surface geometry to cellular alignment, multiscale chirality transfer, and differentiation-associated cell states, highlighting geometry as an important contributor to morphogenetic organization in living matter.

## Materials and Methods

Detailed experimental protocols, materials, substrate-fabrication procedures, image and data analyses, and theoretical derivations are provided in the *SI Appendix*. Briefly, C2C12 mouse and immortalized human myoblasts (IHM) were cultured and differentiated on fibronectin-coated PDMS substrates of defined geometry and curvature, including cylindrical fibers, channels, cones, wavy surfaces, and domes. Fixed and live confocal microscopy were used to quantify cell alignment, migration, differentiation, focal adhesions, and centrosome positioning. Nuclei were segmented with Cellpose (*57*) and tracked with TrackMate (*58*). Actin polymerization, formin activity, and myosin-II activity were perturbed using latrunculin A, SMIFH2, and blebbistatin, respectively. Curvature-dependent organization was interpreted using the continuum nematic framework described in the main text; model fitting and numerical implementation are detailed in the *SI Appendix*.

## Acknowledgments

We thank the members of the “Cell Adhesion and Mechanics” team, Zoheir Guesmia, Léa Trichet and Bruno Cadot for helpful discussions. We thank Dr. Ayako Yamada (ENS-PSL, Paris) for assistance with scanning electron microscope. This work was supported by the European Research Council (Grant No. Adv-101019835 to BL), the Agence Nationale de la Recherche (“Myofuse” ANR-19-CE13-0016 to BL; “STRATEPI” DFG-ANR-22-CE92-0048 to RMM), Institut National du Cancer (INCa_16712 and INCa_18429 to BL and RMM), the “Initiatives d’excellence” (Idex ANR-11-IDEX-0005-02) transverse project BioMechanOE (TP5) (to WX), grant under the 2024 UPCité-NUS call for projects (to WX), the Alexander von Humboldt Foundation (Alexander von Humboldt Professorship to BL), and the Human Frontier Science Program (grant number LT0007/2023-C) (To YS). We acknowledge the ImagoSeine core facility of the IJM, member of IBiSA and France-BioImaging (ANR-10-INBS-04) infrastructures.

## Author contributions

YS, RS, RV, ACJ, and BL conceived and designed the research. YS performed all the experiments with assistance from WX, SD, YLT, JDC, AS, and LA. YS, WX, and GG conducted the microfabrication experiments. YS analyzed the experimental data with input from all authors. RS and ACJ carried out the theoretical modeling and RS carried out the simulations. YS, RS, RV, ACJ, and BL wrote the manuscript, and all authors reviewed and approved the final version.

