## Supplementary Information for "Curvature-guided chiral collective organization of myoblast tissues"

**The file includes the following sections:**

1. Materials and Methods.
2. Supplementary Figures.
3. Legends of Supplementary Movies.
4. Details on data fit in the theoretical model and numerical simulation.
5. References

#### Materials and Methods

*Cell culture and reagents.* C2C12 cells (ATCC CRL-1772) were cultured in DMEM (containing glucose and pyruvate; Life Technologies) supplemented with 10% fetal bovine serum (FBS; Life Technologies) and 1% penicillin-streptomycin (Life Technologies) at 37 ℃ with 5% $CO_2$. IHM cells (*1*) were cultured in DMEM (containing glucose and pyruvate; Life Technologies) supplemented with 20% foetal bovine serum (FBS; Life Technologies), 16% Medium 199 (1X) (containing GlutaMAX; Life Technologies), 0.2 μg $ml^{-1}$ dexamethasone, 0.1 mg $ml^{-1}$ gentamicin (Life Technologies), 5 μg $ml^{-1}$ insulin (human; Sigma-Aldrich), 25 μg $ml^{-1}$ fetuin, 5 ng $ml^{-1}$ hEGF and 0.5 ng $ml^{-1}$ bFGF at 37 ℃ with 5% $CO_2$. In experiments of C2C12 differentiation, cells were cultured in DMEM (containing glucose; Life Technologies) supplemented with 2% horse serum (Life Technologies) and 1% penicillin-streptomycin (Life Technologies) at 37 ℃ with 5% $CO_2$. In experiments of IHM differentiation, cells were cultured in DMEM (containing glucose and pyruvate; Life Technologies) supplemented with 0.1 mg $ml^{-1}$ gentamicin (Life Technologies) and 10 μg $ml^{-1}$ insulin (human; Sigma-Aldrich) at 37 ℃ with 5% $CO_2$. For immunofluorescent staining, cells were fixed with 4% paraformaldehyde (PFA), permeabilized with 0.5% Triton X-100 for 20 min, blocked with 1% BSA/PBS for 1 hour, and incubated with primary antibody overnight at 4 ℃. The samples were then incubated with secondary antibody, phalloidin (1:200; Thermo Fisher) and Hoechst (1:1000; Thermo Fisher) overnight. To characterize cell differentiation, primary antibodies against myosin heavy chain (MHC; DSHB), Pax7 (DSHB), and P27 (BD Biosciences) were used. Cell-substrate adhesions and centrosomes were characterized using primary antibodies against paxillin (Sigma-Aldrich) and pericentrin (Abcam), respectively. To perturb actin polymerization, formin-mediated actin assembly, and myosin II activity, cells were treated with different concentrations of latrunculin A (Sigma-Aldrich), SMIFH2 (Merck), and blebbistatin (Merck), respectively (see main text). To assess drug-induced changes in the intracellular actin cytoskeleton, cells were exposed to the drugs at the time of seeding and fixed after 1 h. To examine their effects on cell motility, cells were cultured in drug-containing medium for 24 h, during which cell movements were recorded by time-lapse microscopy. To investigate their effects on the formation of supracellular helical structures, the drugs were added 1 h after cell seeding, and the cells were subsequently cultured in drug-containing medium for 3 days.

*Microfabrication*. To fabricate PDMS microfibers, freshly mixed silicone elastomer base and curing agent (Sylgard 184, DOWSIL; 10:1, w/w) were first injected into PTFE microtubes (Darwin Microfluidics) of different inner diameters and then incubated at 80 ℃ for 2 hours. After polymerization, the PDMS fibers were pulled out from the PTFE microtubes and cut into pieces of several millimeters in length. The PDMS fibers were then suspended between two glass coverslips of ~300 μm in thickness with a gap of several millimeters. The ends of the fibers were fixed on the coverslips with AB glue. The whole setup was fixed in a glass-bottom petri dish (FluoroDish, catalog no. FD35-100). The fabrication of PDMS channels can be found in our previous work (*2*). Briefly, smooth copper wires (Goodfellow SARL) of different diameters were aligned in parallel several micrometers above a silicon wafer (1 cm by 2 cm) using a homemade stage that could control the positions of each wire precisely. A fresh mixture of silicone elastomer base and silicone elastomer curing agent was poured on the silicon wafer to cover the metal wires. The entire setup was then left at room temperature for several days for PDMS polymerization. After polymerization, the metal wires were pulled out through sonication in acetone. The remaining PDMS block containing straight, parallel microchannels was then cut in a direction perpendicular to the microchannels into small pieces of several millimeters. These small pieces were then stuck to a glass-bottom petri dish through plasma cleaning.

Truncated conical PDMS fibers were fabricated by filling PDMS into sterile pipette tips (eppendorf). The samples were then cured at 80°C for 1 hour. After polymerization, the conical fibers were pulled out from the pipette tips. The conical fibers were then suspended between two glass coverslips of ~300 μm in thickness with a gap of several centimeters. The ends of the fibers were fixed on the coverslips with AB glue. The whole setup was fixed in a glass-bottom petri dish (FluoroDish, catalog no. FD35-100).

Wavy polydimethylsiloxane (PDMS) substrates with controlled wavelengths (Fig. S16A) were fabricated using a glass rod templating technique. Glass rods of defined diameters (Hilgenberg, Germany) were used as templates, with the desired wavelength determined by twice the diameter of the rods. For example, to obtain a wavy substrate with a wavelength of 500 μm, glass rods of 250 μm in diameter were employed. Glass rods (10 cm in length) were cut into ~2 cm segments and aligned in parallel (10–20 rods) on a clean glass slide. A small amount of uncured PDMS was applied to secure the rods in place, and the assembly was cured at 80 °C for 1 hour. After curing, alternating glass rods were carefully removed using fine-tipped tweezers to create a wavy microstructure. Residual PDMS was gently removed, and the surface was thoroughly cleaned by sonication in 100% ethanol for 15 minutes followed by compressed air drying. The resulting wavy template was treated with oxygen plasma for 10 minutes and functionalized by exposure to a silane monolayer (Trichloro (1H, 1H, 2H, 2H-Perfluoroctyl)-silane 97%; Merck) in vapor phase overnight to offer an antifouling surface. The as-fabricated silanized wavy templates were subsequently used for PDMS molding, either to fabricate master molds or to produce wavy substrates for experimental applications.

To fabricate the silicon wafers of the wavy substrates and Gaussian curved surfaces shown in Fig. 1*J-L* and Fig. 5*H-J*, we employed an optimized diffused back-side exposure process (*3*). Briefly, a standard quartz optical mask with chromium-etched dark-field features—comprising lines and circles of varying width, radius, and pitch—was spin-coated with a 250 μm-thick SU-8 negative photoresist. After prebaking at 95 °C for 2 h and allowing natural cooling to room temperature, the resist was exposed to UV LED light (365 nm) using a UV-KUB 2 tool (Kloe, France) for 15 s at an intensity of 30 mW/cm$^2$. The exposure was performed from the back side of the mask through an optical opal diffuser (Edmund Optics, White Diffuser Glass) placed in contact with the mask.

Following exposure, the resist was post-baked on a hot plate at 95 °C for 5 min, developed in SU-8 developer for 10 min, rinsed in fresh IPA, and dried under $N_2$ flow. After hard baking at 120 °C for 2 min, the plate with SU-8 features served as the primary mold.

PDMS replicas were produced using standard soft-lithography: Sylgard 184 PDMS (10:1 base-to-curing-agent ratio) was mixed, degassed, poured onto the primary mold, degassed again in a vacuum jar (10 min at 1–5 mbar), and cured on a hot plate at 60 °C for 2 h. The cured PDMS was peeled off and diced into smaller pieces, each containing a 9 × 9 $mm^2$ pattern. These diced PDMS molds were treated with $O_2$ plasma (20 sccm flow, 2 mbar, 30 W, 30 s) and exposed to the vapor phase of trichloro(1H,1H,2H,2H-perfluorooctyl)silane in a vacuum jar (2–5 mbar) for 1 h. All procedures were performed in an ISO 6 cleanroom.

The PDMS molds were then treated with silanization (Trichloro (1H, 1H, 2H, 2H-Perfluoroctyl)-silane 97%; Merck). After silanization, a fresh mixture of silicone elastomer base and silicone elastomer curing agent was poured on the PDMS molds. The samples were then incubated at 80 ℃ for 2 hours for polymerization. After polymerization, the PDMS substrates with curved microstructures were peeled off from the silicon wafers. All the PDMS substrates were then plasma cleaned for 2 min and treated with silanization (3-Triethoxysilylpropylamine, APTES; Merck). To facilitate cell adhesion, the substrates were coated with a fibronectin solution (50 µg $ml^{-1}$) overnight at 4 ℃.

*Live-cell and fixed-sample imaging.* Live imaging was performed with a 10× objective on a spinning disc microscope (Nikon CSU-W1) at 37 ℃ and 5% $CO_2$. Images were acquired every 20 min in a *z*-stack (2.5 µm per *z* step). To facilitate cell tracking, IHM cells were treated with SPY-DNA/ SPY-FastAct (Tebubio/Spirochrome).

*Data analysis.* Nematic orientational field was analyzed through OrientationJ, a plugin of ImageJ. The nematic order parameter in Fig. 2*A* is defined as $S_n = \sqrt{\langle \cos 2\alpha \rangle^2 + \langle \sin 2\alpha \rangle^2}$ where $\alpha$ is the angle between $\boldsymbol{n}$ and the *x*-axis, $\langle \ldots \rangle$ represents the average over the whole field of view. To track the motion of individual cells, cell nuclei were first segmented by Cellpose-2.0 (*4, 5*) and then tracked by Trackmate (a plugin of ImageJ) (*6*). The velocity order parameter of cell motion shown in Fig. 2*A* is defined as $S_v = \sqrt{\langle \cos 2\beta \rangle^2 + \langle \sin 2\beta \rangle^2}$, where $\beta$ is the angle between cell velocity and the *x*-axis. The spatial velocity correlation function is defined as $C_v(\delta r) = \frac{\langle \boldsymbol{v}_i(0)\cdot \boldsymbol{v}_j(\delta r)\rangle}{\langle \boldsymbol{v}_i(0)\cdot \boldsymbol{v}_i(0)\rangle}$, where $\boldsymbol{v}$ represents the velocity vector of cells, $\delta r$ represents the distance between cell pairs, $\langle \ldots \rangle$ represents average over different cell pairs *i* and *j*. Similarly, the temporal velocity correlation function is defined as $C_v(\delta t) = \frac{\langle \boldsymbol{v}_i(0)\cdot \boldsymbol{v}_i(\delta t)\rangle}{\langle \boldsymbol{v}_i(0)\cdot \boldsymbol{v}_i(0)\rangle}$, where $\delta t$ represents time delay between the velocity vectors of the same cell, $\langle \ldots \rangle$ represents average over different cells *i*. The inertia tensor of trajectory structures is defined according to refs. (*7, 8*). In brief $M_{ab} = \sum_{i=1}^{N} (\rho_a^i - \overline{\rho_a})(\rho_b^i - \overline{\rho_b})$, where *N* represents the total number of trajectory points, *a*, *b* represents *x*, *y*; $\boldsymbol{\rho}^i = (\rho_x^i, \rho_y^i)$ are coordinates of the *i*th trajectory point; and $\overline{\boldsymbol{\rho}} = N^{-1} \sum_{i=1}^{N} \boldsymbol{\rho}^i$ is the center of mass of the structure formed by the trajectory points. This allows us to define the anisotropy of the structure, $q = \frac{(M_{xx} - M_{yy} + 2iM_{xy})}{(M_{yy} + M_{xx})} = |q| e^{i2\phi}$, where $|q|$ and $\phi$ are the structure's elongation and the azimuthal angle of its long axis, respectively. The segmentations of cell nuclei in Fig. 6*B* were completed with Cellpose-2.0 (*4, 5*).

Centrosome-based analysis of single-cell chirality within monolayers. To quantify single-cell chirality within dense monolayers, we analyzed the orientation of the nucleus-centrosome axis relative to the local nematic director. Nuclei and centrosomes were identified from fluorescence images, and for each cell the nucleus-centrosome axis was defined as the vector pointing from the nuclear centroid to the centrosome position (Fig. S15). The orientation angle of this axis was denoted as $\alpha$, with $-\pi < \alpha \leq \pi$. The local nematic director around each cell was obtained from the actin orientation field using OrientationJ, and its orientation angle was denoted as $\beta$. Because the local cell alignment is nematic, $\beta$ and $\beta + \pi$ are equivalent, i.e., $-\pi/2 < \beta \leq \pi/2$. We therefore calculated the signed angular difference between the nucleus-centrosome axis and the local nematic director as:

$$\delta = \frac{1}{2} \times \mathrm{atan2}\{\sin[2 \times (\alpha - \beta)], \cos[2 \times (\alpha - \beta)]\},$$

where atan2($a$,$b$) calculates the angle between the vector ($b$, $a$) and the positive $x$-axis. This maps $\delta$ into the interval $(-\pi/2, \pi/2]$. With this convention, $\delta > 0$ corresponds to a counterclockwise rotation of the nucleus-centrosome axis relative to the local nematic director, whereas $\delta < 0$ corresponds to a clockwise rotation.

Temporal velocity fluctuations. To calculate the temporal velocity fluctuations of cells on fibers of diameter 300 and 800 μm, we first track the motion of individual cells with Cellpose and Trackmate. We then calculate the velocity of each cell as a function of time, $\mathbf{v}_i(t)$ as well as its longitudinal (along the long-axis of fiber) and transverse (perpendicular to the long-axis of fiber) components, $v_x^i(t)$ and $v_y^i(t)$. For each cell, we then get a time series of speeds from which we calculate their standard deviation, $\delta v_i = \sqrt{\frac{1}{T_i}\sum_{t=1}^{T_i}(v_i(t) - \mu_i)^2}$, where $T_i$ represents the number of time points of cell $i$, $v_i(t)$ represents the speed of cell $i$ at time $t$, and $\mu_i$ represents the mean speed of cell $i$ over time $T$. Similar for the fluctuations of longitudinal, $\delta v_x^i$, and transverse, $\delta v_y^i$, components.

Spatial velocity fluctuations. To measure the spatial speed fluctuations of cells, we calculate the standard deviations of the speed, $\delta v(t)$, of different cells on fibers at each time moment, $\delta v(t) = \sqrt{\frac{1}{N(t)}\sum_{i=1}^{N(t)}(v_i(t) - \mu(t))^2}$, where $N(t)$ are the total cell number at time $t$, $v_i(t)$ represents the speed of cell $i$ at time $t$, and $\mu(t) = \frac{1}{N(t)}\sum_{i=1}^{N(t)} v_i(t)$ represents the mean cell velocity at time $t$, and then plot the standard deviation as a function of time. Similar for the spatial fluctuations of the longitudinal, $\delta v_x(t)$, and transverse, $\delta v_y(t)$, components of velocities.

#### Supplementary Figures

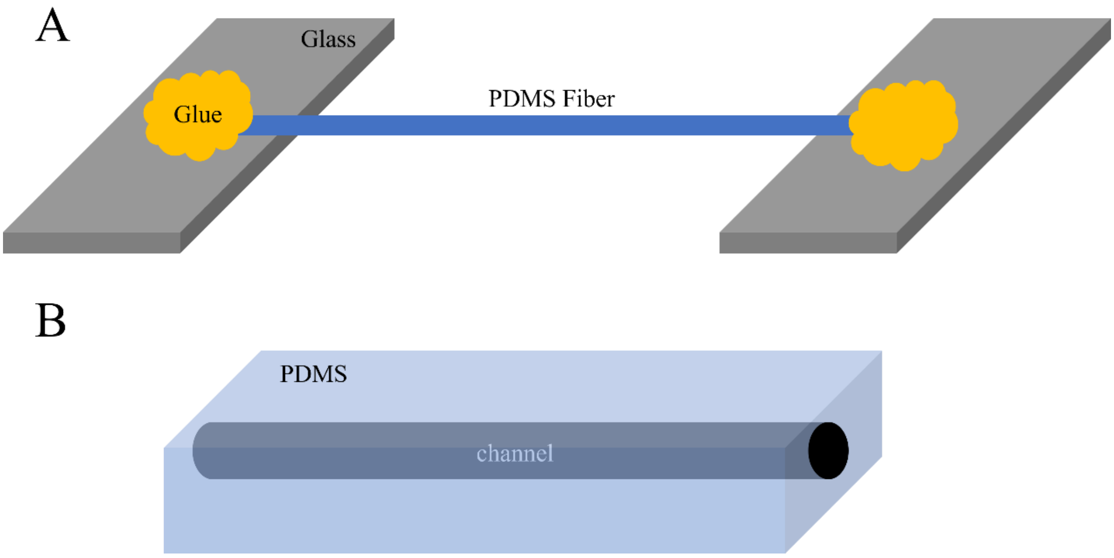


**Fig. S1** (A) Schematic structure of a PDMS fiber suspended on two pieces of glass substrates and fixed by glue at the two ends. (B) Schematic structure of a PDMS channel.

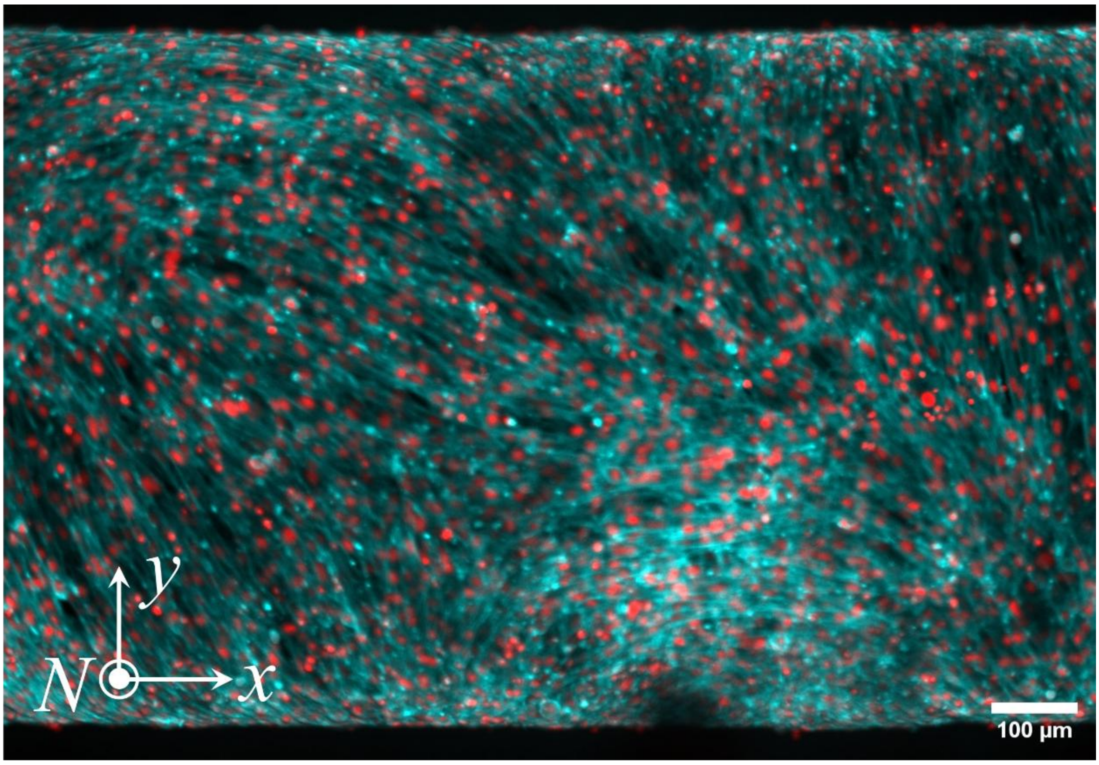


**Fig. S2** A typical example showing the degenerate alignment of C2C12 cells on a fiber of diameter $D \approx 800$ μm. Red: nuclei, Cyan: actin.

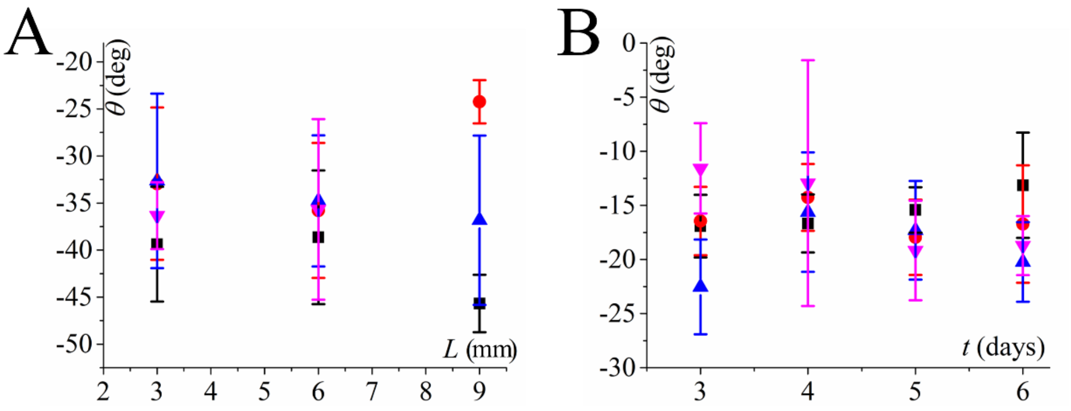


**Fig. S3** (A) The dependence of the tilt angle ($\theta$) of C2C12 cell on the length of fibers ($L$), fiber diameter $D$=500 μm. (B) The dependence of the tilt angle ($\theta$) of C2C12 cell on the culture time ($t$), fiber diameter $D$=300 μm. Each data point represents a single fiber. The error bars represent the standard deviation of the tilt angle of the cells on the fiber.

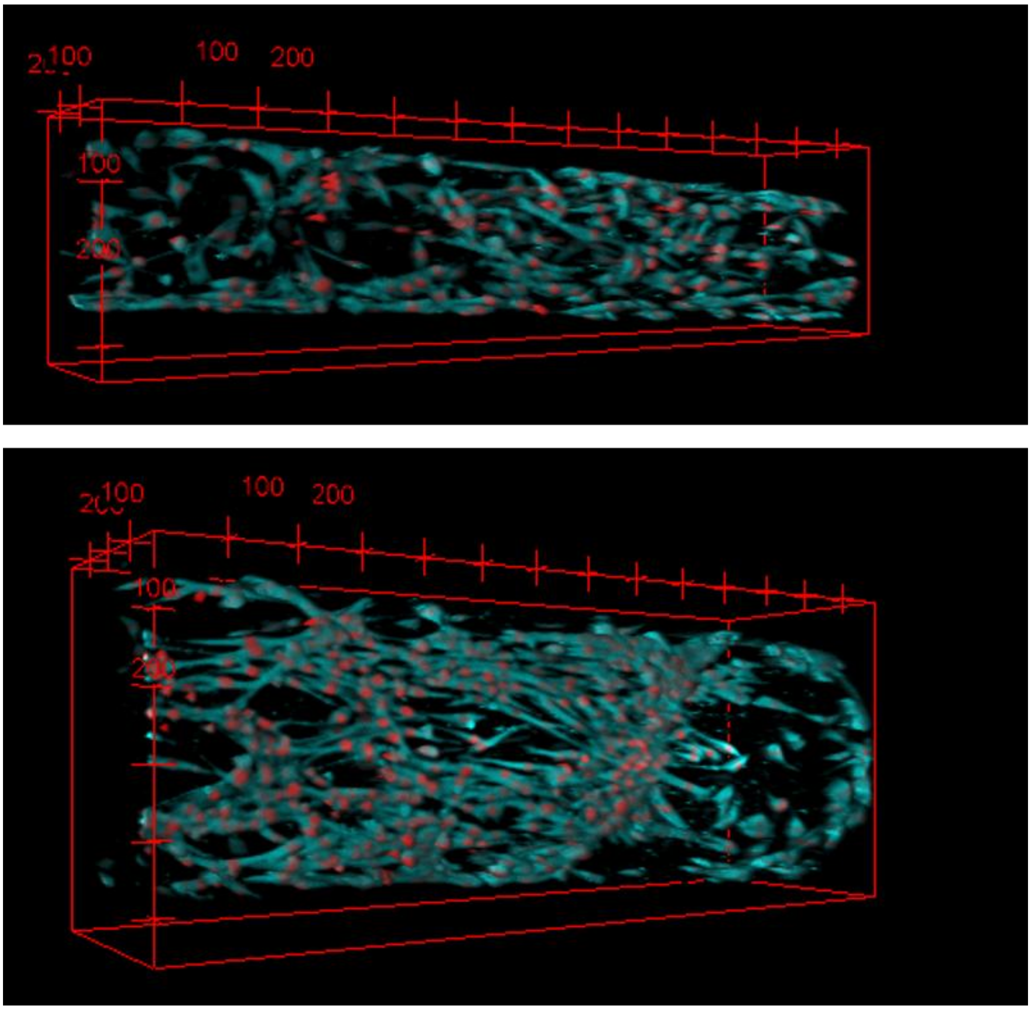


**Fig. S4** 3D fluorescent microscopic images showing that C2C12 myoblasts fail to spread out and form helical superstructures in channels (top $D$ = 200 μm, bottom $D$ = 500 μm).

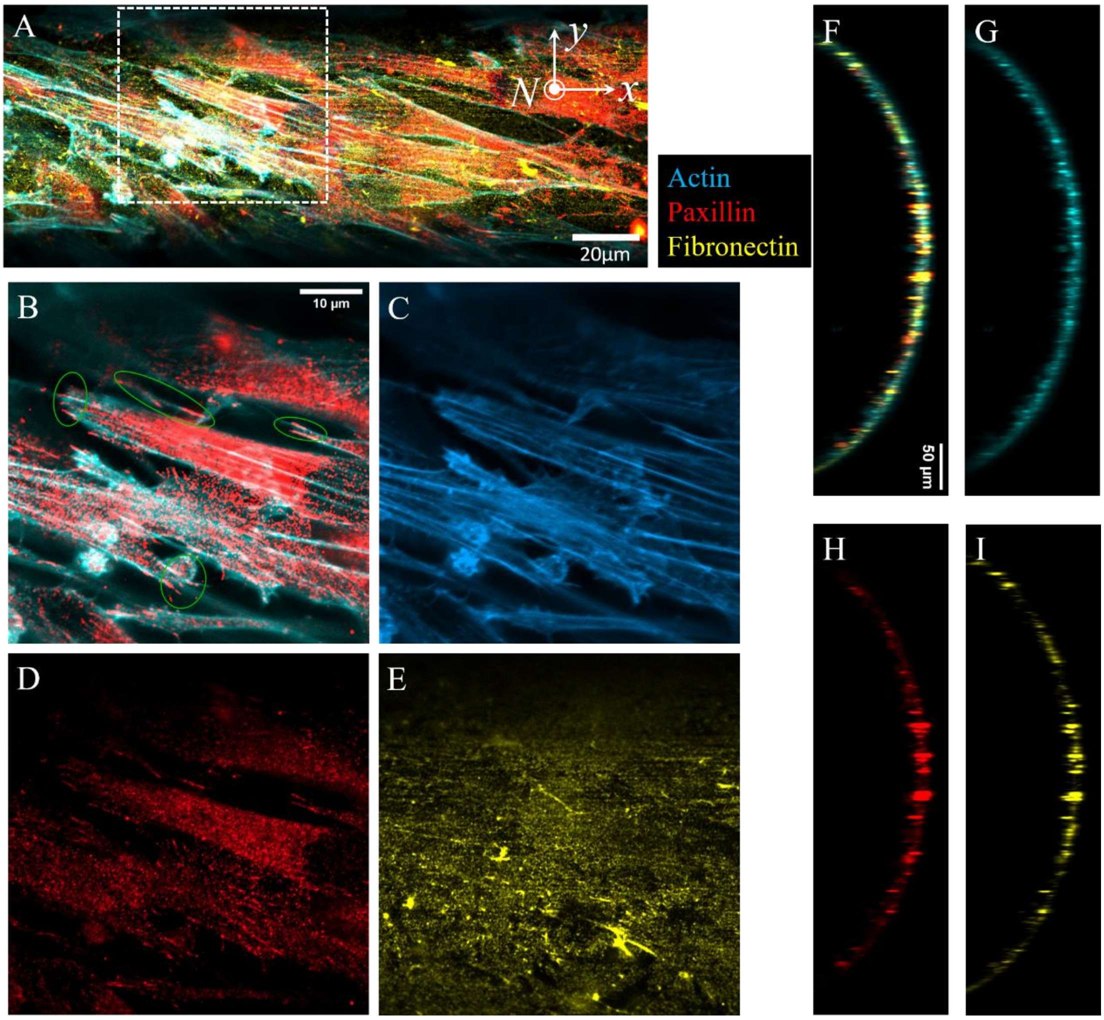


**Fig. S5** Cell-matrix adhesion in channels. (A) The *z*-stack projection of the confocal images of C2C12 cells in a channel of diameter of 500 µm. The cyan, red and yellow colors represent cell actin, cell paxillin and the coated fibronectin, respectively. (B) The magnified field of view showing the paxillin and actin of cells corresponding to the region of the white-dashed square in (A). The green circles mark the focal adhesions of the cells. (C-E) The corresponding different channels of (B). (F) The *z*-stack projection of the cross-sectional confocal image shows that cells remain closely attached to the fibronectin coated curved substrate. (G-I) The corresponding different channels of (F).

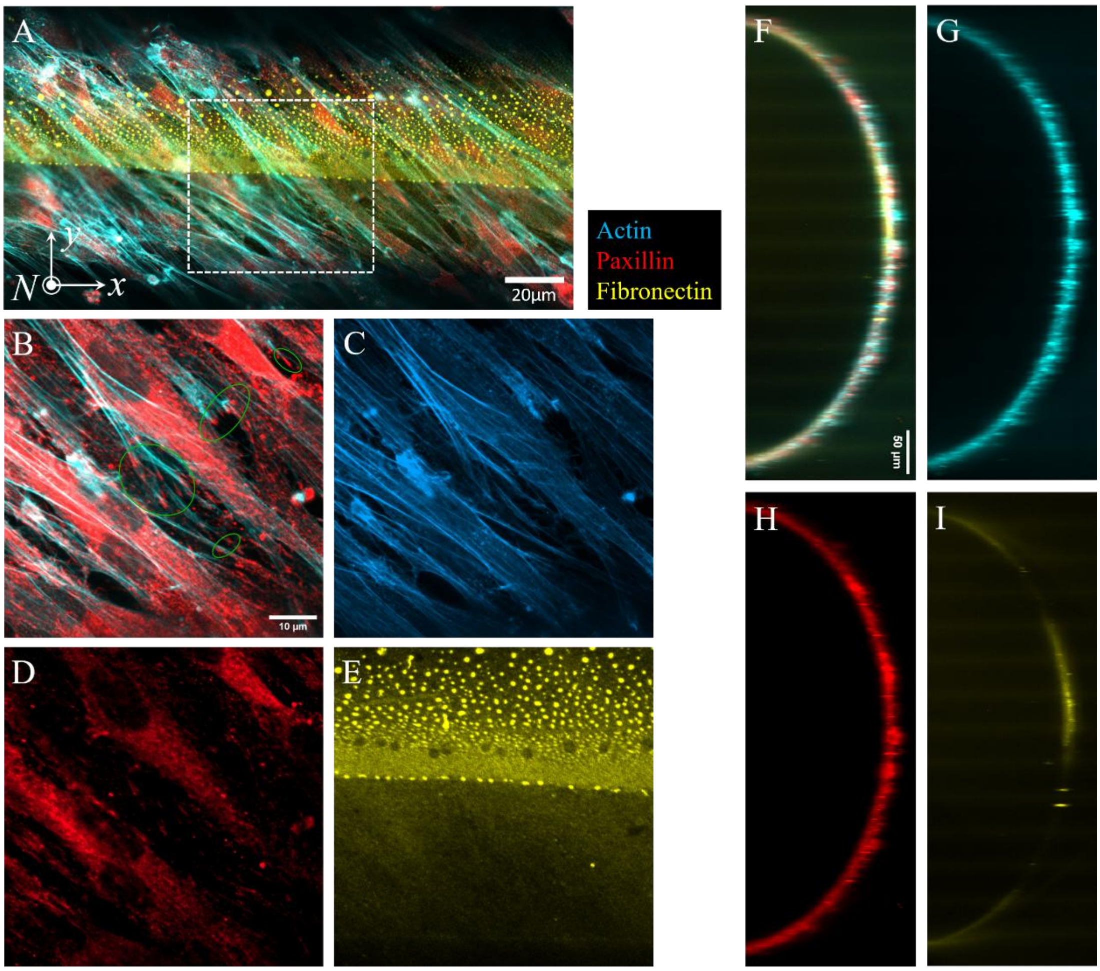


**Fig. S6** Cell-matrix adhesion on fibers. (A) The *z*-stack projection of the confocal images of C2C12 cells on a fiber of diameter of 500 μm. The cyan, red and yellow colors represent cell actin, cell paxillin and the coated fibronectin, respectively. (B) The magnified field of view showing the paxillin and actin of cells corresponding to the region of the white-dashed square in (A). The green circles mark the focal adhesions of the cells. (C-E) The corresponding different channels of (B). (F) The *z*-stack projection of the cross-sectional confocal image shows that cells remain closely attached to the fibronectin coated curved substrate. (G-I) The corresponding different channels of (F).

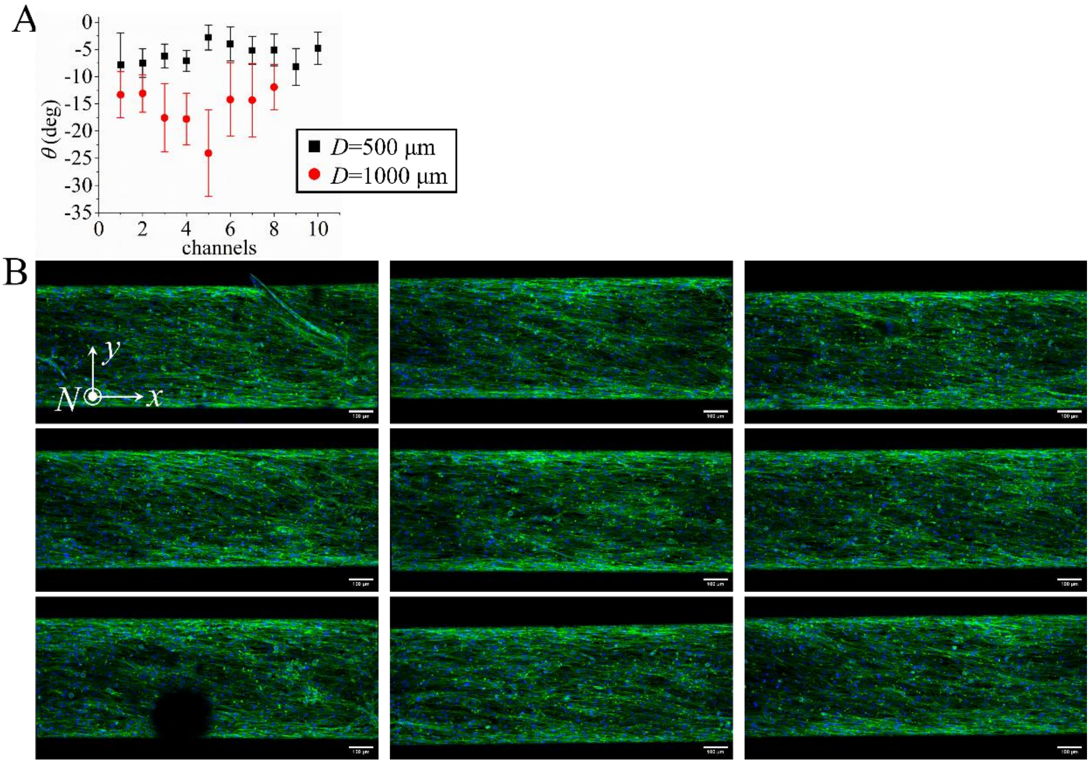


**Fig. S7** Alignment of C2C12 in channels. (A) The mean values of the tilted angle $\theta$ of the director field of C2C12 in different channels (black: $D$=500μm; red: $D$=1000μm). Each data point represents a single channel. The error bars represent the standard deviation of the tilt angle of the cells in the channel. (B) Typical examples showing the fluorescent microscopic images of C2C12 cells on the bottom section of different channels of $D$=500μm. Green: actin. Blue: nucleus. Scale bars 100 μm.

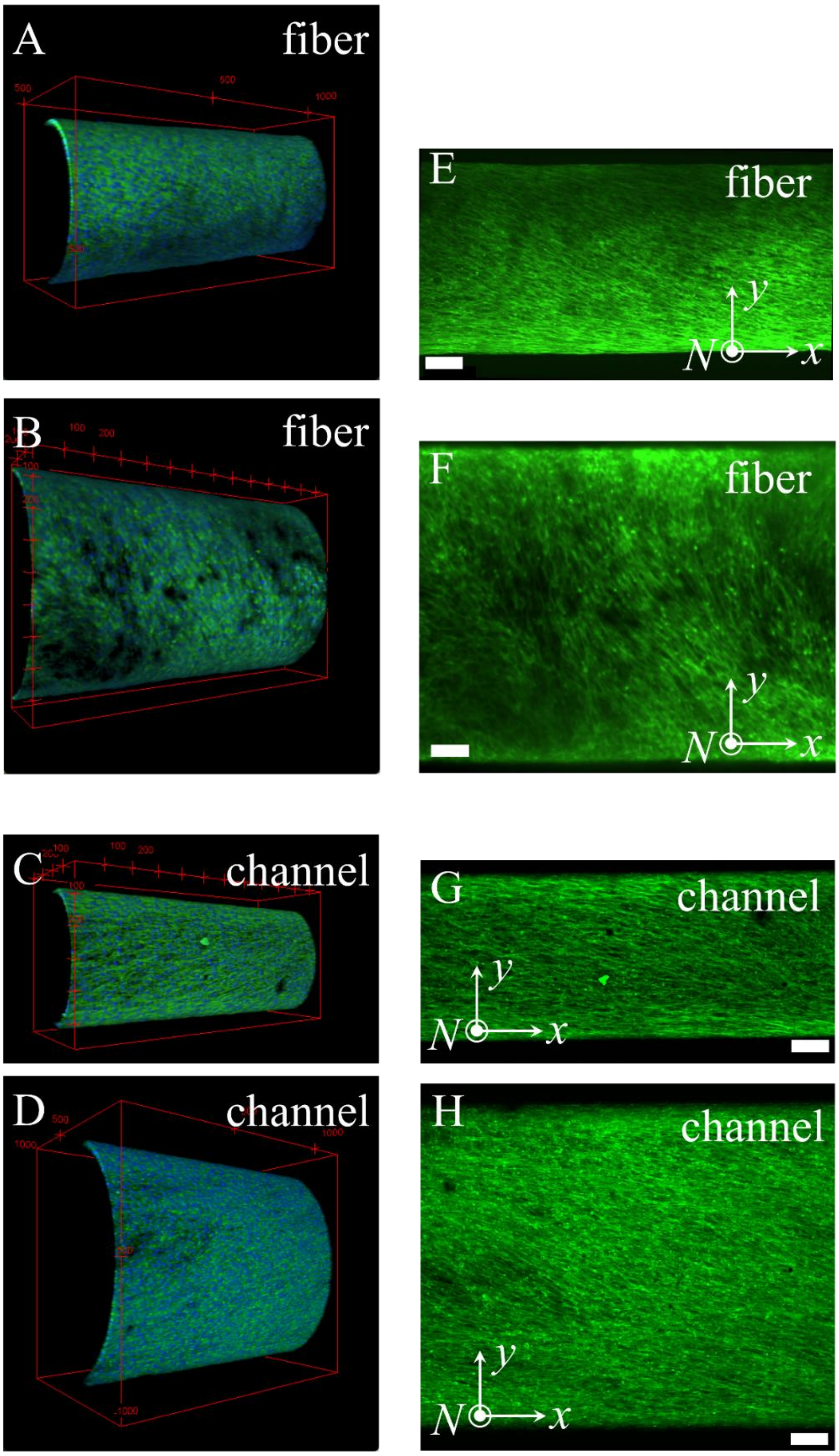


**Fig. S8** IHM on fibers and channels of different diameters. (A) 3D fluorescent microscopic image of IHM on a fiber with $D \sim 500$ µm. (B) 3D fluorescent microscopic image of IHM on a fiber with $D \sim 800$ µm. (C) 3D fluorescent microscopic image of IHM in a channel with $D \sim 500$ µm. (D) 3D fluorescent microscopic image of IHM in a channel with $D \sim 1000$ µm. (E) and (F) show the *z*-stack projection of the myoblasts in (A) and (B). (G) and (H) show the *z*-stack projection of the myoblasts in (C) and (D). Scale bars 100 µm. Green: actin; Blue: nucleus.

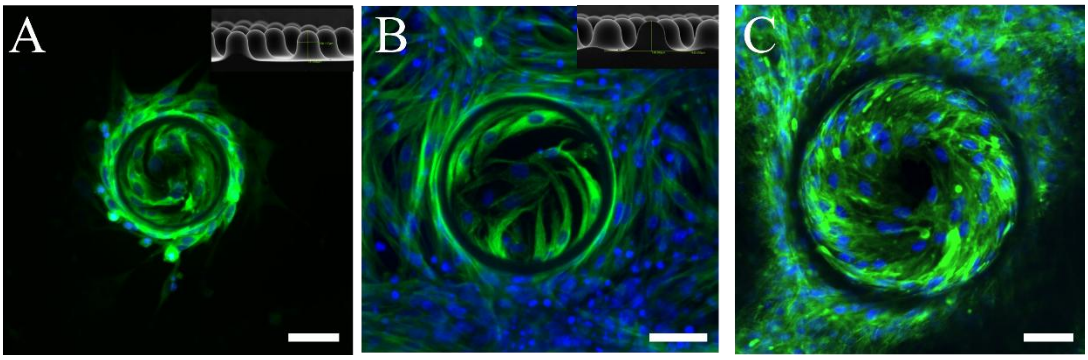


**Fig. S9** Fluorescent microscopy images (green: actin, blue: nuclei) of IHM on different domes. (A) and (B) The *z*-stack projections of fluorescent microscopic images of IHM on Gaussian-like domes (apical view). The insets show the corresponding SEM images of the substrates. Scale bars 50 μm. (C) The *z*-stack projection of the fluorescent microscopic images of IHM on a half spherical dome (apical view). Scale bar 50 μm.

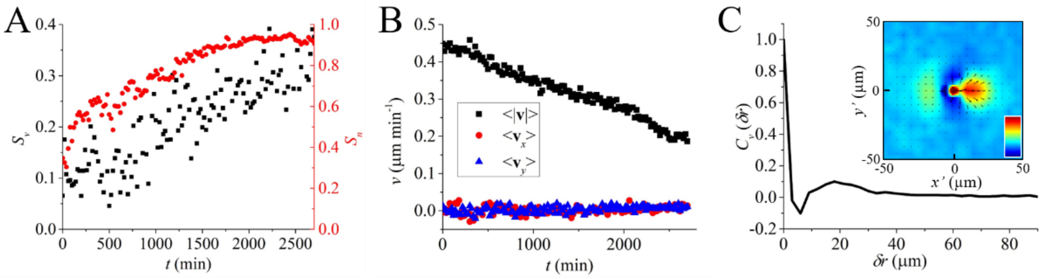


**Fig. S10** Dynamics of IHM on a fiber. (*A*) Temporal dependence of velocity order parameter ($S_v$, black squares) and nematic order parameter ($S_n$, red circles), respectively. (*B*) Temporal dependence of cell velocity. Black squares represent the mean value of cell speed. Red circles and blue triangles represent the mean values of the components of cell velocity along the *x*- and *y*-axes, respectively. (*C*) 1D spatial velocity correlation function of IHM on the fiber. Inset: 2D spatial velocity correlation function of IHM on the fiber. The color bar scales linearly from -0.2 (dark blue) to 0.5 (dark red).

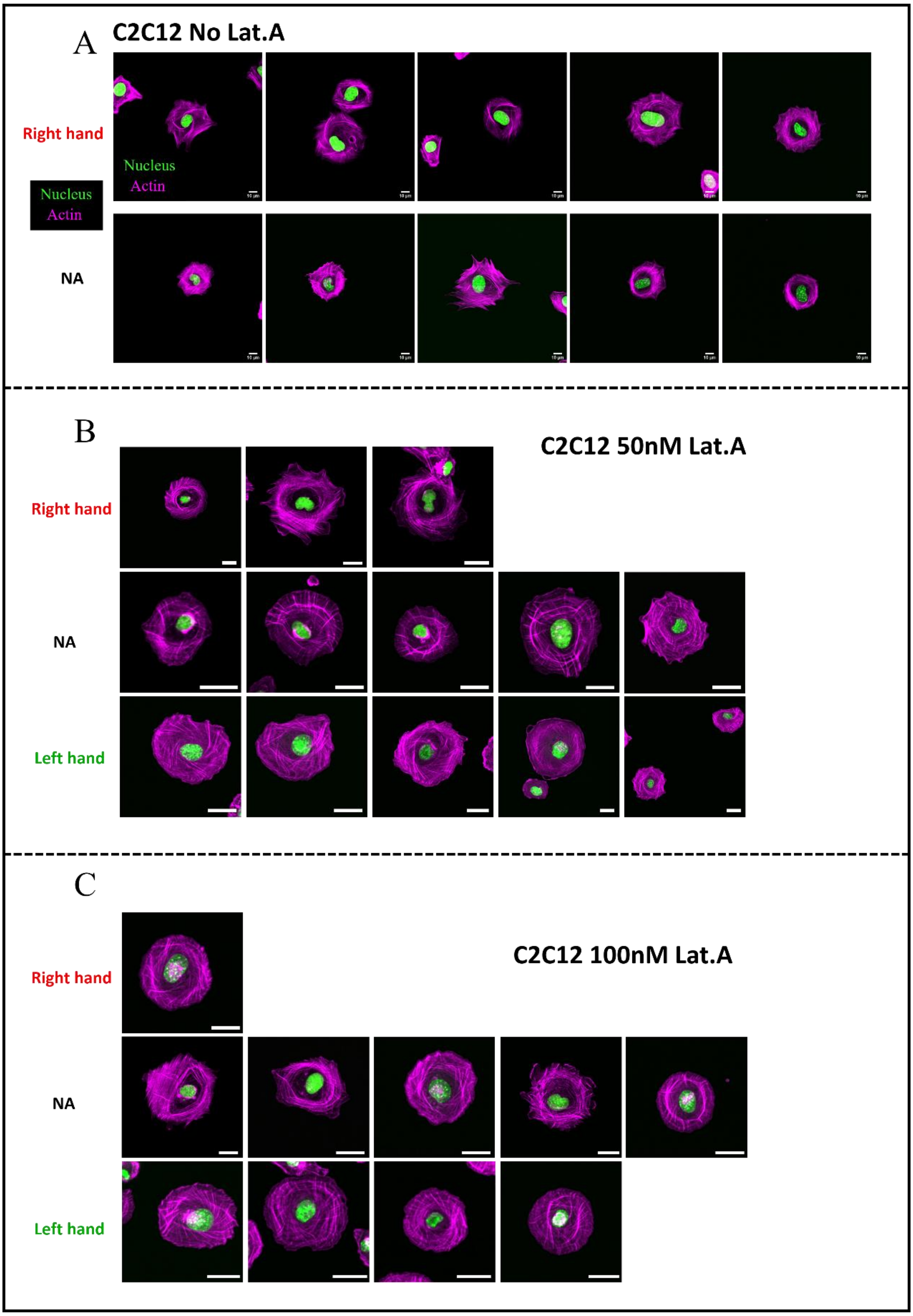


**Fig. S11** Typical examples show the spiral actin pattern in C2C12 cells. The C2C12 cells in (B) and (C) are treated with 50nM and 100nM Latrunculin-A, respectively. Right hand represents cells shows spiral actin pattern with right-handed chirality; Left hand represents cells shows spiral actin pattern with left-handed chirality; NA represents cell which do not show clear spiral actin pattern. Scale bars are 10 µm in (A) and 20 µm in (B) and (C).

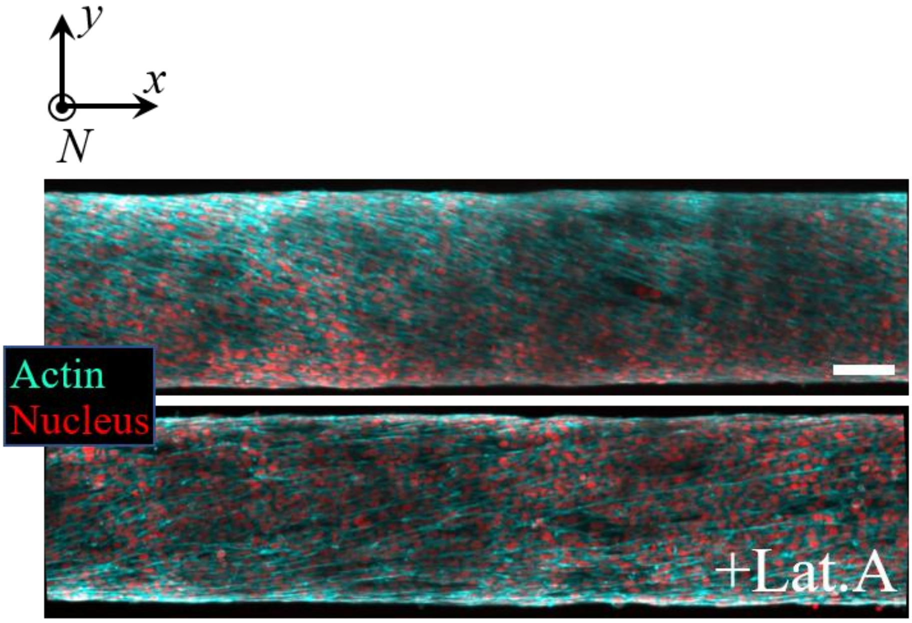


**Fig. S12** *Z*-stack projections of fluorescent microscopic images of helical superstructures of IHM on fibers processed with (bottom) and without (top) *Lat-A* treatment, respectively. Scale bar 100 μm.

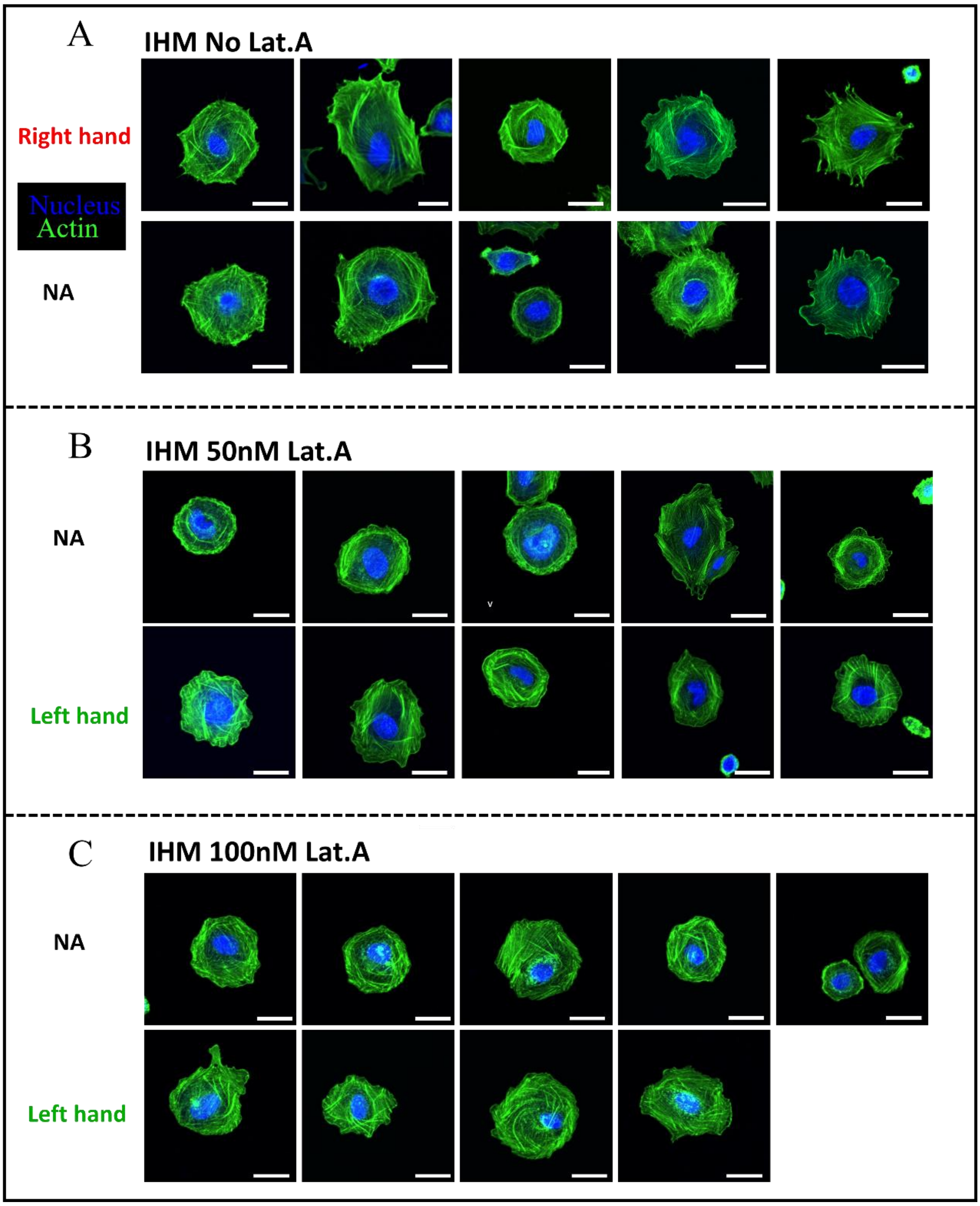


**Fig. S13** Typical examples show the spiral actin pattern in IHM cells. The IHM cells in (B) and (C) are treated with 50nM and 100nM Latrunculin-A, respectively. Right hand represents cells shows spiral actin pattern with right-handed chirality; Left hand represents cells shows spiral actin pattern with left-handed chirality; NA represents cell which do not show clear spiral actin pattern. Scale bars 20 μm.

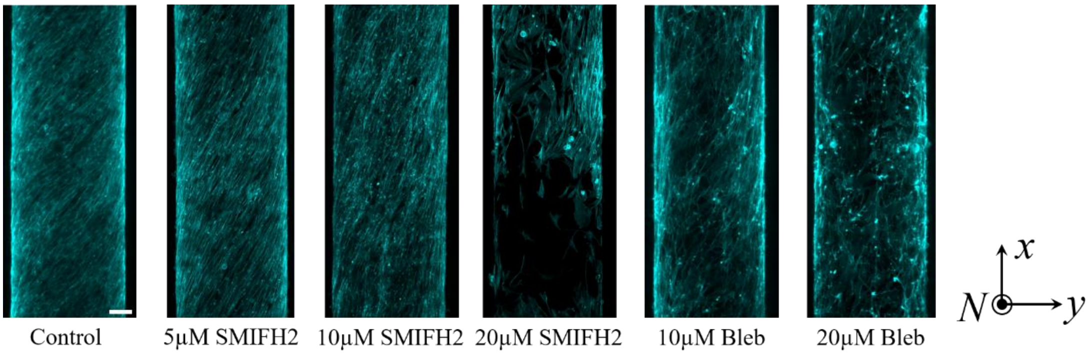


**Fig. S14** The *z*-stack projection of confocal images of the actin of C2C12 cells on fibers at different conditions. Scale bar 100 μm.

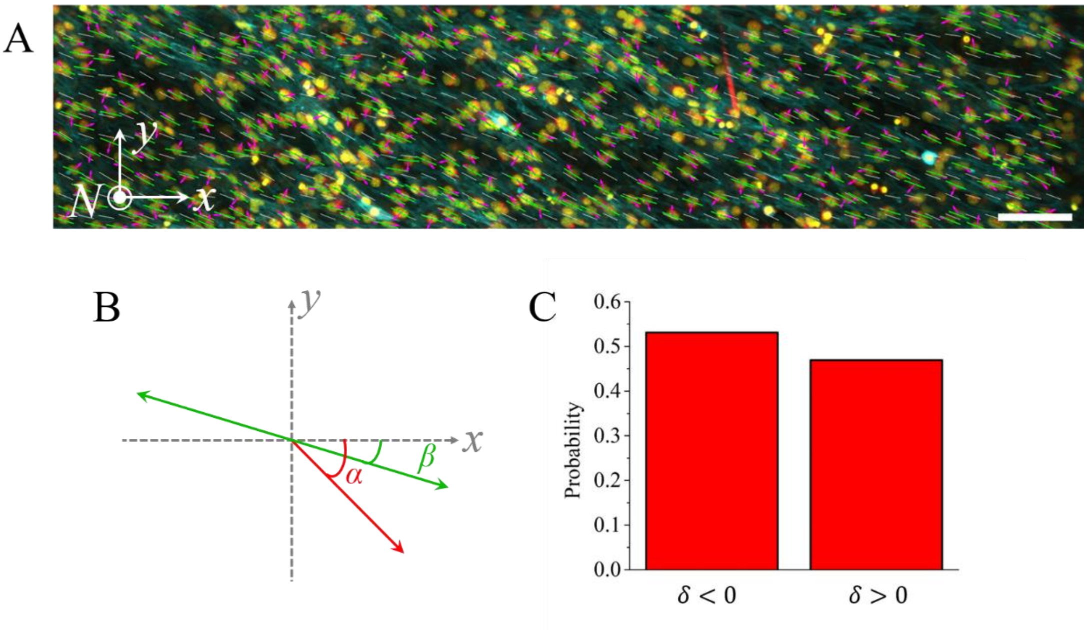


**Fig. S15** Cell chirality in dense layers. (A) z-stack projection of confocal images of C2C12 cells on a fiber. The cyan, yellow and red colors represent cell actin, nucleus and centrosome, respectively. The white-short lines represent the nematic director field obtained from OrientationJ. The green short lines represent the local nematic director averaged over the nematic director field within a small window (50μm × 50μm) around the reference cell. The magenta arrow represent the vector from nucleus centroid to centrosome. (B) A schematic drawing shows the local nematic director (green arrow) and the nucleus-centrosome axis (red arrow) and their angles $\beta$ and $\alpha$ with respect to the positive $x$-axis direction. (C) Statistics showing the probability of positive and negative $\delta$. $\delta > 0$ corresponds to a counterclockwise rotation of the nucleus-centrosome axis relative to the local nematic director, whereas $\delta < 0$ corresponds to a clockwise rotation. More than 1000 cells from two independent samples are measured.

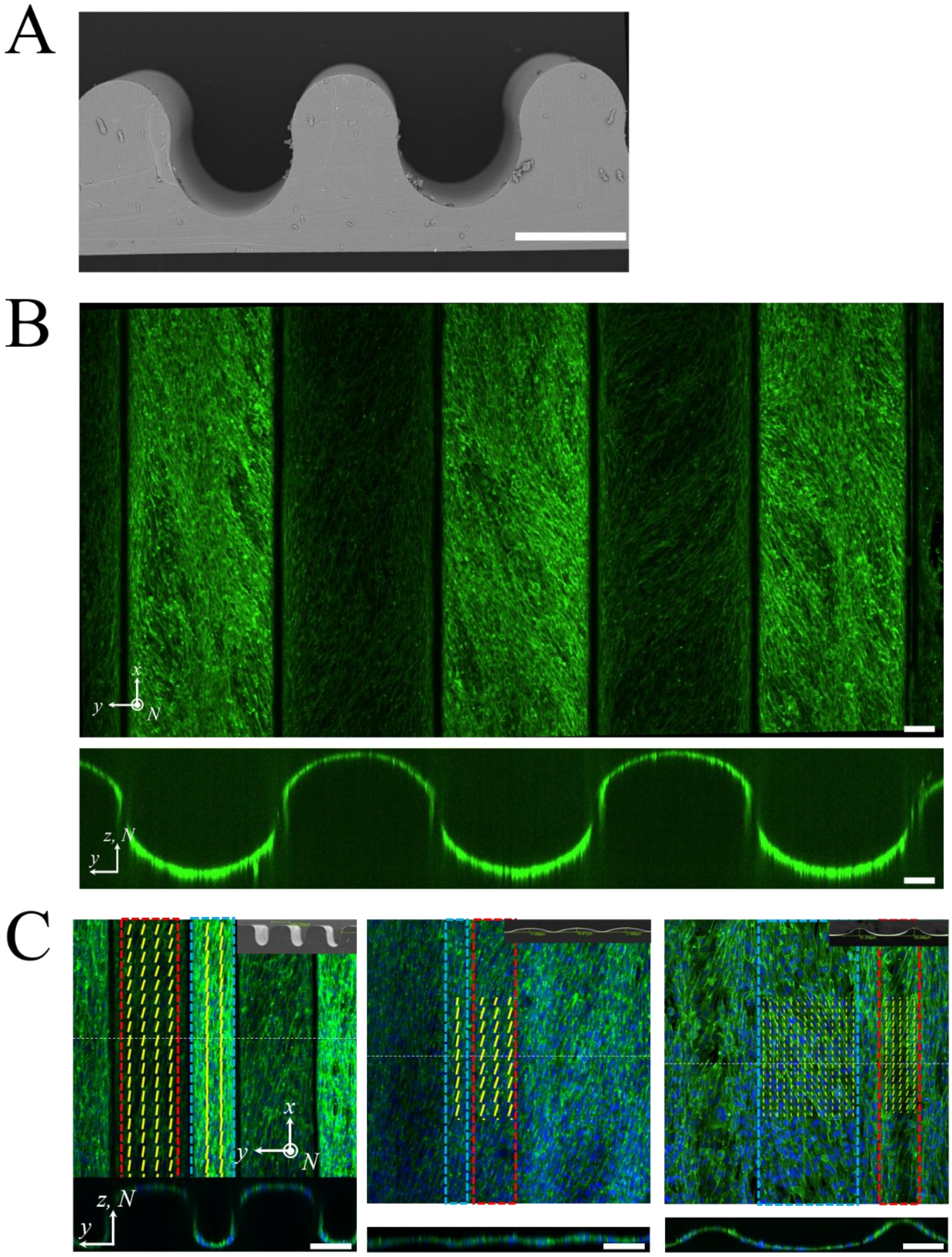


**Fig. S16** (A) Scanning electron microscopy (SEM) of the wavy substrate. Scale bar 500 μm. (B) IHM on a wavy substrate. The top image shows the *z*-stack projection of fluorescent microscopic images of IHM on a wavy substrate (apical view). The bottom image shows the cross section of the sample in the *yz*-plane. Scale bars 100 μm. (C) Fluorescent microscopy images (green: actin, blue: nuclei) of IHM on different wavy surfaces. The *z*-stack projections (top) and the cross sections (bottom) of the fluorescent microscopic images of IHM on different wavy substrates (apical view). The red and blue dashed rectangles represent the ridges and valleys, respectively. The insets show the corresponding scanning electron microscopy (SEM) images of the PDMS substrates. The yellow dashed lines represent the director field obtained by OrientationJ. Scale bars 100 μm.

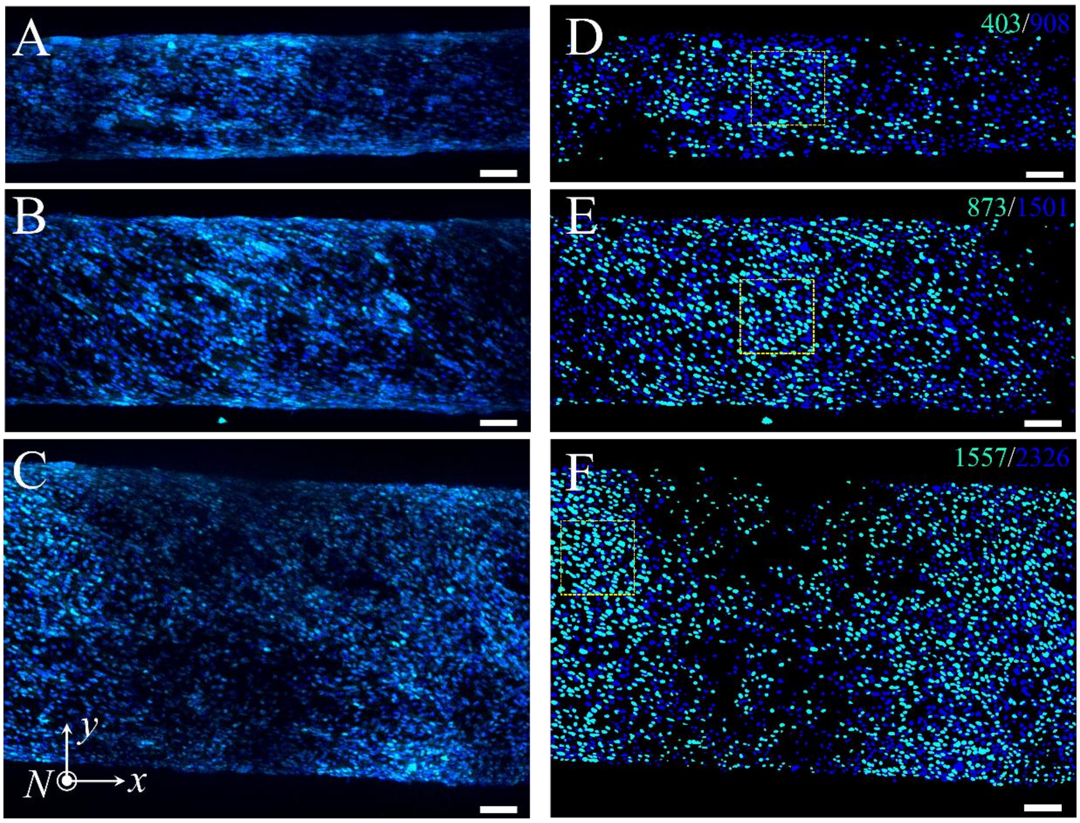


**Fig. S17** (A)-(C) *Z*-stack projection of fluorescent microscopy images of IHM cells cultured on fibers of different diameters, stained for nuclei (blue, DAPI) and p27 (cyan). (D)-(F) The corresponding nuclei segmentation images processed by Cellpose. The regions in yellow dashed squares represent the images shown in Fig. 6B. The cyan and blue numbers in (D)-(F) give the number of p27-positive cells and total number of cells in the image. Scale bars 100 μm.

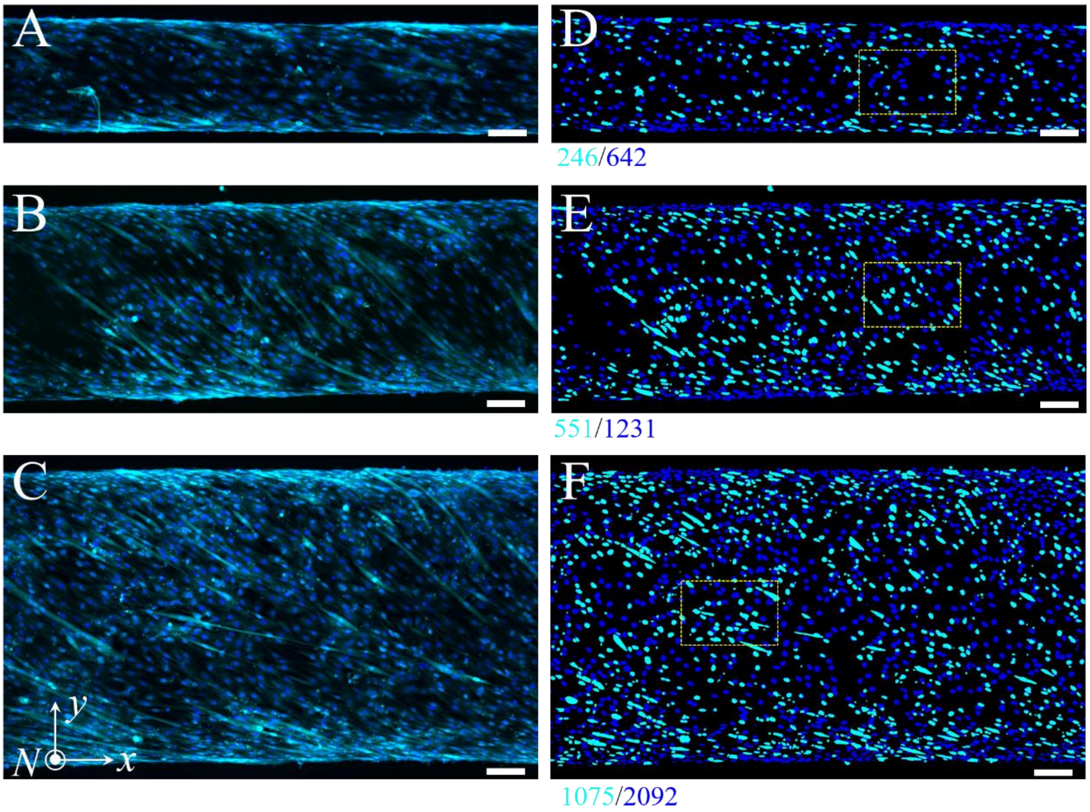


**Fig. S18** (A)-(C) *Z*-stack projection of fluorescent microscopy images of C2C12 cells cultured on fibers of different diameters, stained for nuclei (blue, DAPI) and p27 (cyan). (D)-(F) The corresponding nuclei segmentation images processed by Cellpose. The regions in yellow dashed squares represent the images shown in Fig. 6B. The cyan and blue numbers in (D)-(F) give the number of p27-positive cells and total number of cells in the image. Scale bars 100 μm.

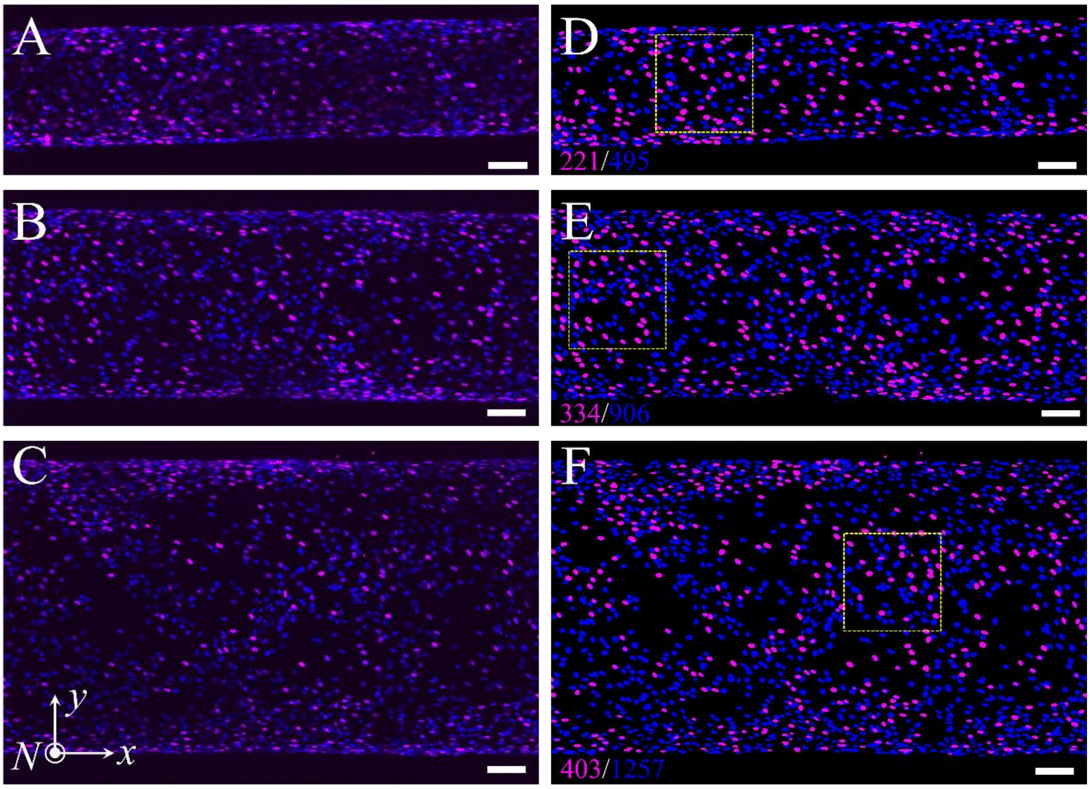


**Fig. S19** (A)-(C) *Z*-stack projection of fluorescent microscopy images of C2C12 cells cultured on fibers of different diameters, stained for nuclei (blue, DAPI) and Pax7 (magenta). (D)-(F) The corresponding nuclei segmentation images processed by Cellpose. The regions in yellow dashed squares represent the images shown in Fig. 6B. The magenta and blue numbers in (D)-(F) give the number of Pax7-positive cells and total number of cells in the image. Scale bars 100 μm.

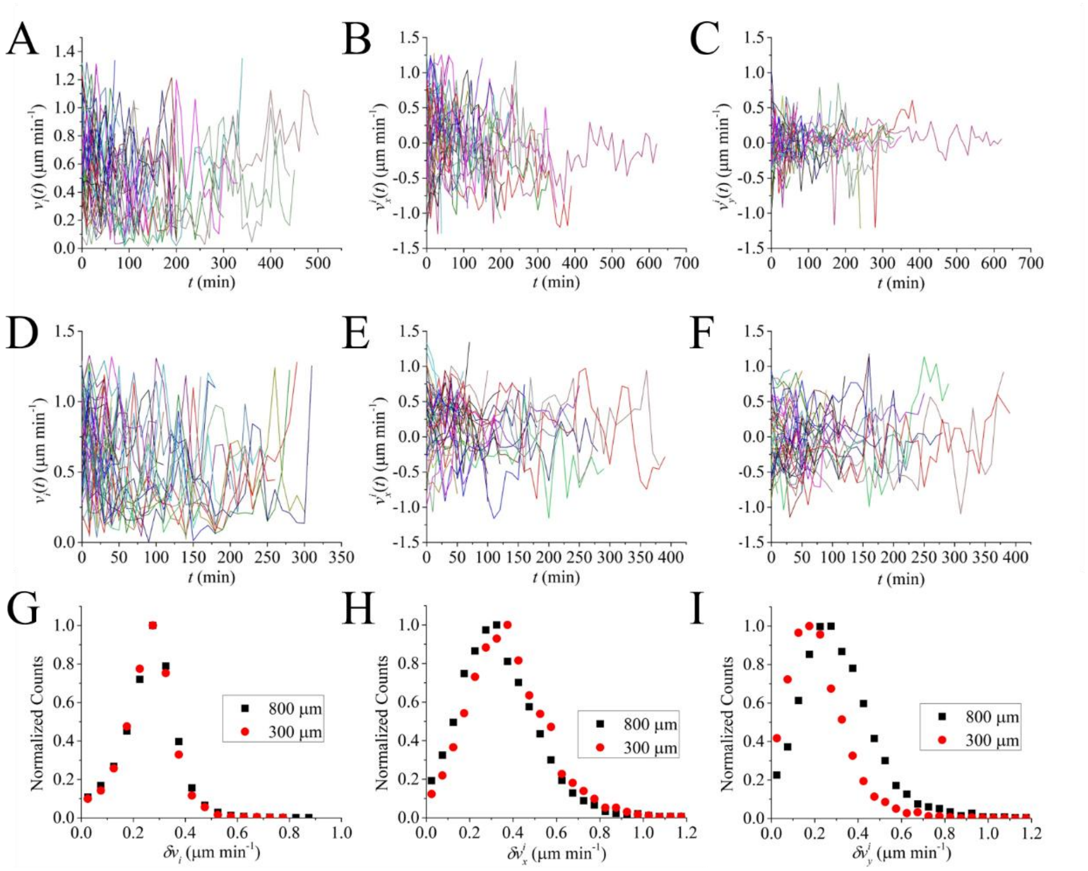


**Fig. S20** Temporal velocity fluctuation of cells on fibers. (A-C) show the speed ($v_i$), longitudinal ($v_x^i$) and transverse ($v_y^i$) components of the velocities of 50 randomly selected cells on a fiber whose diameter is 300 μm as a function of time. (D-F) show the speed ($v_i$), longitudinal ($v_x^i$) and transverse ($v_y^i$) components of the velocities of 50 randomly selected cells on a fiber whose diameter is 800 μm as a function of time. (G-I) show the normalized probability distribution of the temporal fluctuations of the cell speed (G) and the longitudinal (H) and transverse (I) components of cell velocities.

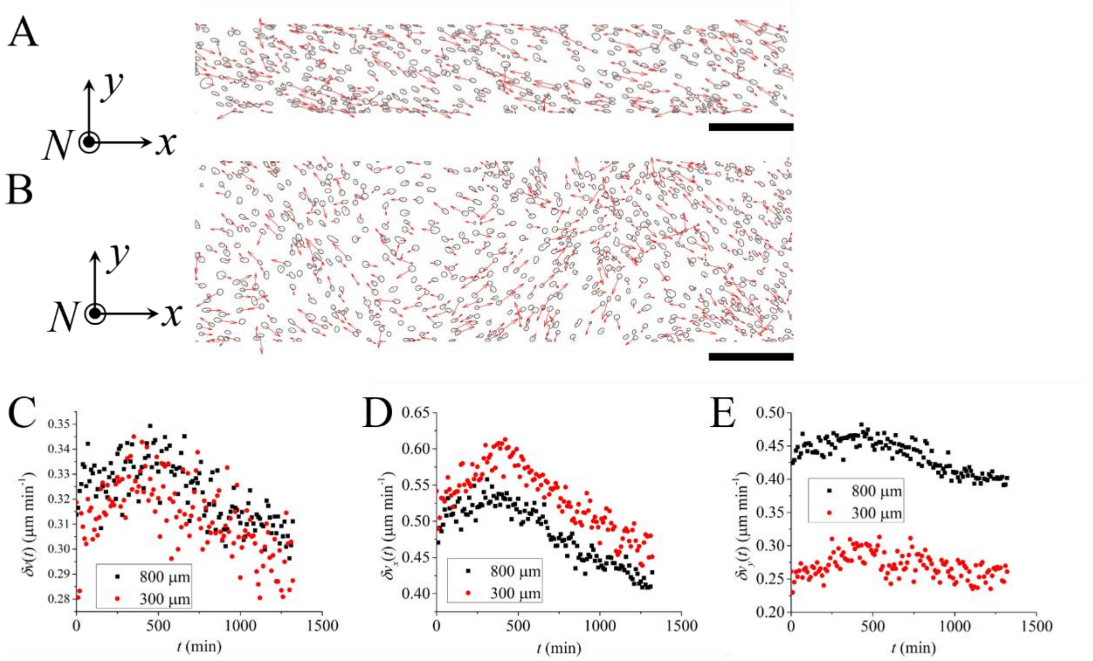


**Fig. S21** Spatial velocity fluctuations of cells on fibers. (A-B) shows a snapshot of cell movement on fibers of diameters of 300 μm (A) and 800 μm (B), respectively. Scale bars 200 μm. (C-E) spatial fluctuations of cell speed ($\delta v(t)$, C), longitudinal velocity component ($\delta v_x(t)$, D) and transverse velocity component ($\delta v_y(t)$, E) as a function of time.

#### Supplementary Movies

**Movie S1.** The formation process of the helical supracellular structure of C2C12 cells on a fiber of diameter 300 μm. Green: nucleus; Red: actin.

**Movie S2.** The dynamics of the spiral actin organization in an isolated cell.

**Movie S3.** The dynamics of actin filaments in cells within a dense monolayer. The cell marked by the white square shows transient actin flows from cell periphery to center at the very beginning of the movie, but the actin flows are soon disturbed by the interactions of neighboring cells.

**Movie S4.** The dynamics of nuclei of C2C12 cells in a channel.

**Movie S5.** The dynamics of nuclei of C2C12 cells in a valley.

### Supplementary Theory

We provide here some supporting derivations and details to accompany the main text. In Sec. 1 we provide details on the procedure used to obtain the parameters underlying the fit in Fig. 3K. In Sec. 2 we describe the method used to simulate active chiral nematic order on a truncated cone (Fig. 4F).

### 1 Details on fit procedure leading to Fig. 3K

Here, we briefly discuss the fitting procedure that we have used to fit our model, Eq. 3 of the main text, to the experimental data; see Fig. 3K.

To fit our model, we used the following cost function:

$$\mathcal{L}(h, \boldsymbol{\eta}) = \sum_i \sum_j \left\{ w_i \left(\mathrm{Re}[\theta(C_i, \boldsymbol{\eta})] - \theta_{ij}\right)^2 + \mathrm{Im}[\theta(C_i, \boldsymbol{\eta})]^2 \right\} \mathbf{1}_{L[\theta(C_i,h)]>0} \,. \tag{S1}$$

For clarity, we recall Eq. 3 of the main text, which defines the model prediction

$$(k_{\mathrm{bend}} C^2 + k_{\mathrm{ud}} C) \sin(2\theta) + k_{\mathrm{chi}} C \cos(2\theta) - \gamma\Omega = 0. \tag{S2}$$

We denote the three parameters in the model—namely $\gamma\Omega/k_{\mathrm{bend}}$, $k_{\mathrm{chi}}/k_{\mathrm{bend}}$, and $k_{\mathrm{ud}}/k_{\mathrm{bend}}$—collectively as a parameter vector $\boldsymbol{\eta}$. Furthermore, $\theta_{ij}$ are the experimental readings at curvature $C_i$, where multiple independent measurements are indexed by $j$. The sets of data are the measured nematic tilt angles on fibers and channels (Figs. 1f and S7A). Here $\mathrm{Im}(\cdot)$ and $\mathrm{Re}(\cdot)$ denote the imaginary and real parts, respectively.

The first term in $\mathcal{L}$ is a standard least-squares term. If, for a given $\boldsymbol{\eta}$, the model prediction $\theta(C_i, \boldsymbol{\eta})$ at curvature $C_i$ is real, then this corresponds to a uniform, steady-state solution to Eq. 1 of the main text. In contrast, a complex $\theta(C_i, \boldsymbol{\eta})$ would signal a non-steady state solution. This can occur, for example, if the active rotation $\Omega$ is too large to be balanced by restoring torques deriving from the free energy. If this were the case, then we would not observe uniform steady-states, which is not the case for the range of curvatures explored in our experiments. We therefore include the second term in curly braces to penalize complex solutions. Also, we consider weighted least-squares penalty, where the weight $w_i = {\sigma_i^{-2}}/\frac{1}{N}\sum_k {\sigma_k^{-2}}$.

Finally, we note that for given curvature $C$ and parameters $\boldsymbol{\eta}$, Eq. 3 of the main text yields multiple solutions for $\theta$, one of which is stable. To exclude the unstable solutions, we include the indicator function $\mathbf{1}_{L[\theta(C_i,h)]>0}$ in the definition of $\mathcal{L}$, where

$$L[\theta(C_i, \boldsymbol{\eta})] = \left[ (k_{\mathrm{bend}} C_i^2 + k_{\mathrm{ud}} C_i) \cos(2\theta) - k_{\mathrm{chi}} \sin(2\theta) \right] / \gamma\Omega \,. \tag{S3}$$

and require

$$L[\theta(C_i, \boldsymbol{\eta})] > 0.$$

This expression is nothing more than $\partial^2 f/\partial\theta^2$, where $f$ is the Frank free energy density (Eq. 2 of the main text). Thus, $L > 0$ corresponds to a linearly stable steady state.

We use the bootstrap method to estimate the parameters [9]. In each bootstrap iteration, the experimental observations for each diameter $D$ were resampled with replacement. For these sampled values, the parameter vector $\boldsymbol{\eta}$ was estimated by minimizing the loss function using a stochastic gradient algorithm. By applying bootstrapping, we obtained the distribution of $\boldsymbol{\eta}$, for which we report in the main text the mean as well as the 95% confidence intervals, defined by the 2.5th and 97.5th percentiles. The shaded bands in Fig. 3K denote the corresponding 95% bootstrap prediction intervals.

### 2 Details on numerical simulation leading to Fig. 4F

To model the coexistence of ordered and disordered monolayer regions observed experimentally on the conical surface (Fig. 3) we developed a numerical code for the dynamics of the nematic order parameter tensor, $\mathbf{Q}$. We assume that these dynamics, in the absence of fluid motion, can be modeled with an active Landau–de Gennes approach, consisting of relaxational model-A dynamics governed by an energy $\mathcal{F} = \int dS\, f_Q(\mathbf{Q}, \nabla\mathbf{Q}, \mathbf{C})$, supplemented by a non-variational active torque, due to substrate-mediated driving. We note that we allow for a generic dependence of $f_Q$ on substrate whose curvature is described by the second-rank tensor $\mathbf{C}$. We thus write

$$\partial_t \mathbf{Q} + \Omega\,(\boldsymbol{\epsilon}\cdot\mathbf{Q} - \mathbf{Q}\cdot\boldsymbol{\epsilon}) = -\frac{1}{\Gamma}\frac{\delta\mathcal{F}}{\delta\mathbf{Q}}\,. \tag{S4}$$

In this equation $\Omega$, oriented normally to the surface, is the rate of rotation arising from active torques; $\boldsymbol{\epsilon}$ is the two-dimensional Levi-Civita tensor; and $\Gamma$ is a rotational viscosity associated with $\mathbf{Q}$.

The free energy $\mathcal{F}$ is taken here to be a generalization of the one that we employed using the director approximation ($S_n \approx 1$); see Eq. 2 in the main text. That is, $\mathcal{F} = \mathcal{F}_{\rm LdG} + \mathcal{F}'$. In this decomposition, $\mathcal{F}_{\rm LdG}$ is the standard Landau (bulk) and de Gennes (gradient) terms in the one-constant approximation:

$$\mathcal{F}_{\rm LdG} = \int d\mathcal{S}\left[\frac{a}{2}\,\mathrm{Tr}(\mathbf{Q}^2) + \frac{c}{4}\left(\mathrm{Tr}(\mathbf{Q}^2)\right)^2 + \frac{\mathcal{K}}{2}\,|\nabla\mathbf{Q}|^2\right], \tag{S5}$$

where $a < 0$, $c > 0$, and $\mathcal{K} > 0$ are phenomenological coefficients. We also note that $|\nabla\mathbf{Q}|^2 = \nabla_i Q_{jk}\nabla^i Q^{jk}$, where $\nabla_i$ is a covariant derivative; please see Ref. [10] for our differential geometric notation. The second part of the total energy contains contributions from anisotropic bending energy; and from broken up-down and chiral symmetries:

$$\mathcal{F}' = \int d\mathcal{S}\left[K_{\rm bend}\,\mathrm{Tr}(\mathbf{C}^2\cdot\mathbf{Q}) + K_{\rm ud}\,\mathrm{Tr}(\mathbf{C}\cdot\mathbf{Q}) + K_{\rm chi}\,\mathrm{Tr}(\boldsymbol{\epsilon}\cdot\mathbf{C}\cdot\mathbf{Q})\right], \tag{S6}$$

where $K_{\rm bend} > 0$ and the other coefficients may be of either sign.

For numerical purposes it is convenient to write Eq. S4 in terms of a matrix representation of $\mathbf{Q}$. Note that the position vector of a point on the conical surface can be written $\mathbf{X} = r(s)\hat{\mathbf{r}} + z(s)\hat{\mathbf{z}}$ where $s$ is the arc length along one of its generators, $r(s)$ is it radial position and $z(s)$ its axial position. The tangent basis is then

$$\mathbf{e}_s = r'(s)\hat{\mathbf{r}} + z'(s)\hat{\mathbf{z}} \tag{S7}$$

$$\mathbf{e}_\phi = r(s)\hat{\boldsymbol{\phi}}\,, \tag{S8}$$

where $\phi$ is the azimuthal angle. The components of $\mathbf{Q}$, related to the local director $\mathbf{n}$ and the degree of alignment $S_n$ via

$$\mathbf{Q} = S_n(\mathbf{n}\otimes\mathbf{n} - \mathbf{I}/2)\,, \tag{S9}$$

are then given by $Q_{ij} = \mathbf{e}_i\cdot\mathbf{Q}\cdot\mathbf{e}_j$:

$$q_1 \equiv Q_{11} = \frac{S_n}{2}\cos 2\theta \tag{S10}$$

$$q_2 \equiv \frac{Q_{12}}{r} = \frac{S_n}{2}\sin 2\theta\,, \tag{S11}$$

where $\theta$ is the tilt angle. Thus, when $\mathbf{Q}$ is expressed in terms of $q_1$ and $q_2$, after some algebra one obtains from Eq. S4 the following coupled equations:

$$\partial_t q_1 + 2\,\Omega\,q_2 = \frac{\mathcal{K}}{\Gamma}\left(\nabla^2 q_1 - \frac{4\,r'^2}{r^2}\,q_1 - \frac{4\,r'}{r^2}\,\partial_\phi q_2\right) - \frac{a}{\Gamma}\,q_1 - \frac{2c}{\Gamma}\left(q_1^2+q_2^2\right)q_1 - \frac{K_{\rm ud}}{\Gamma}\,\mathcal{D}_1 - \frac{K_{\rm bend}}{\Gamma}\,\mathcal{D}_2 \tag{S12}$$

$$\partial_t q_2 - 2\,\Omega\,q_1 = \frac{\mathcal{K}}{\Gamma}\left(\nabla^2 q_2 - \frac{4\,r'^2}{r^2}\,q_2 + \frac{4\,r'}{r^2}\,\partial_\phi q_1\right) - \frac{a}{\Gamma}\,q_2 - \frac{2c}{\Gamma}\left(q_1^2+q_2^2\right)q_2 + \frac{K_{\rm chi}}{\Gamma}\,\mathcal{D}_1\,. \tag{S13}$$

In the above the Laplacian of $q_\alpha$, $\alpha = 1, 2$, is

$$\nabla^2 q_\alpha = \partial_s^2 q_\alpha + \frac{r'}{r}\,\partial_s q_\alpha + \frac{1}{r^2}\,\partial_\phi^2 \tag{S14}$$

and

$$\mathcal{D}_1 = C_s{}^s - C_\phi{}^\phi \tag{S15}$$

$$\mathcal{D}_2 = \left(C_s{}^s\right)^2 - \left(C_\phi{}^\phi\right)^2 \tag{S16}$$

are the curvature terms that couple to bending and to up-down/chiral energies, respectively.

Finally, a short note on the numerics used in solving Eqs. S12 and S13. The initial conditions correspond to a disordered (isotropic) state. We apply periodic boundary conditions along $\phi$ and along $z$ (by gluing the truncated cone with its mirror image to form a “bowtie”, and then periodically repeating this domain). We chose the following parameters, which are consistent with the fit parameters obtained from Figs. 3K and 4E: $\Gamma\Omega/K_{\rm bend} \sim \gamma\Omega/k_{\rm bend} = -10^{-4}$ $\mu$m$^2$; $K_{\rm chi}/K_{\rm bend} \sim k_{\rm chi}/k_{\rm bend} = -10^{-3}$ $\mu$m$^{-1}$; $K/K_{\rm bend} = 0.07$; and $K_{\rm ud}/K_{\rm bend} \sim k_{\rm ud}/k_{\rm bend} = -10^{-3}$ $\mu$m$^{-1}$; please see Ref. [11] for the general correspondence between parameters in the director and tensor descriptions. With these parameters, the coupled equations were solved numerically using the Dedalus package (we used IVP solver with SBDF2 time integrator scheme) [12].
